# Deregulation of multiple mechanisms shapes the onset of *LAMA2*-congenital muscular dystrophy

**DOI:** 10.1101/2024.01.20.576409

**Authors:** Susana G Martins, Vanessa Ribeiro, Catarina Melo, Cláudia Paulino-Cavaco, Dario Antonini, Sharadha Dayalan Naidu, Fernanda Murtinheira, Inês Fonseca, Bérénice Saget, Mafalda Pita, Diogo R Fernandes, Pedro G dos Santos, Gabriela Rodrigues, Rita Zilhão, Federico Herrera, Albena T Dinkova-Kostova, Ana Rita Carlos, Sólveig Thorsteinsdóttir

## Abstract

*LAMA2*-congenital muscular dystrophy (LAMA2-CMD) is the most common congenital muscular dystrophy. This often-lethal disease is triggered by mutations in *LAMA2*, coding for laminin-α2 chain, a key extracellular matrix (ECM) component, prevalent in the skeletal muscle. Several phenotypes have been associated with LAMA2-CMD, however, it is not yet known what mechanisms are faulty, right at disease onset *in utero*. Using the *dy^W^* mouse model of LAMA2-CMD we showed that the disease onset is characterized by a profound downregulation of gene expression, with a marked effect on cytoskeletal organization, myoblast differentiation and fusion and altered DNA repair and oxidative stress responses. Concordantly, we found that *Lama2*-deficient myoblast cells displayed proliferation and differentiation defects, increased oxidative stress and DNA damage. Together, our findings provide unique insights into the processes dependent on laminin-α2 chain during muscle development, revealing its critical importance to maintain muscle cell homeostasis already at fetal stages.

## INTRODUCTION

Muscle formation and growth are processes that start early in embryonic development and continue postnatally. In a first wave, during primary myogenesis, muscle stem cells (MuSCs) dissociate from the dermamyotome and migrate to their target sites, marking the areas where the future muscle will develop^1^. The MuSCs (PAX3+/PAX7+) initiate their differentiation program, which is marked by the activation of the myogenic regulatory factors (MRFs) MYF5, MYOD, MRF4 and myogenin, and give rise to primary myoblasts and finally to the primary (or embryonic) myotubes (myosin heavy chain expressing cells)^1,2^. While some MuSCs differentiate, others remain stem cells (PAX7+) and proliferate substantially until they receive a second stimuli to start differentiating. Differentiation and fusion of these MuSCs lead to the formation of secondary myotubes, marking the beginning of the secondary (or fetal) myogenesis. The transition between primary and secondary myogenesis is promoted by the transcription factor nuclear factor one X (NFIX), which activates fetal-specific genes and blocks embryonic genes^3,4^. Myogenesis is therefore a tightly regulated process involving the contribution of distinct players including transcription factors, growth factor-induced signaling pathways, and extracellular matrix (ECM)^5–7^.

The ECM is a dynamic and vital non-cellular structure within tissues that offers continuous support by forming a scaffold that facilitates the maintenance of tissue morphogenesis and homeostasis. Moreover, the ECM, and particularly the basement membrane (*i.e.* a specialized type of ECM), is essential to assure a correct communication between the extracellular and intracellular environments and the activation of signaling cascades important for cell function^8,9^. The ECM consists of a network of glycoproteins (e.g. laminins, fibronectin), collagens, proteoglycans and other factors, and its composition and function differ according to the tissue and the stage of development^8,9^. In fact, ECM remodeling is part of normal tissue development, and is important for homeostasis and regeneration. During embryonic development, the ECM is constantly changing as tissues and organs are formed and new functions are acquired^8^.

In skeletal muscle, the ECM plays a critical role regulating muscle development, growth and repair, and is vital for effective muscle contraction and force transmission ^1,7,10^. Not surprisingly, mutations in genes encoding ECM components have been associated with a set of conditions collectively termed muscular dystrophies, which are characterized by progressive muscle weakness and loss of muscle mass. One such condition is *LAMA2*-congenital muscular dystrophy (LAMA2-CMD), which is caused by mutations in the *LAMA2* gene, codifying for the α2 chain of laminins 211 (the most abundant laminin isoform in the basement membrane of adult skeletal muscle) and 221^11,12^. LAMA2-CMD is a devastating and often fatal neuromuscular disease, in which patients display hypotonia from birth. Studies using post-natal mouse models of the disease and biopsies from LAMA2-CMD patient have identified several mechanisms linked to the pathology of the disease, including proliferation and differentiation defects, as well as increased oxidative stress and metabolic alterations^11–18^. Additionally, patients suffering from this pathology develop chronic inflammation and fibrosis, which are considered to be secondary events that arise from constant ECM remodeling and cycles of degeneration and regeneration of the muscle fibers^11,12^. To date, the majority of studies aimed at unraveling the mechanisms underlying this disease have been performed postnatally, after the secondary events have arisen. However, our prior data using the *dy^W^* mouse model for LAMA2-CMD indicates that the disease onset occurs during fetal development, specifically between embryonic days (E) 17.5 and E18.5^19^. It is characterized by a decrease in the number of MuSCs/myoblast and impaired muscle growth, even though the muscle appears morphologically normal^19^. In this study, we designed both *in vitro* and *in vivo* approaches to dissect the molecular and cellular processes that fail and are at the core of LAMA2-CMD onset. Our approach revealed that the absence of laminin-α2 chain impairs cell proliferation and differentiation and triggers oxidative stress and DNA damage. RNA sequencing analysis revealed a massive downregulation of gene expression in muscle fibers at E17.5, supporting defective muscle differentiation and fusion and alterations in the cytoskeleton. Our work raises the possibility that the absence of *Lama2* disrupts intra- and extracellular signaling and cell biomechanics, compromising muscle structure and function. The identification of these profound perturbations already at fetal stages, when the muscle appears morphologically normal and before secondary events set in, indicates that they are part of the primary disease mechanism.

## RESULTS

### *Lama2*-deficiency leads to proliferation and cell cycle defects *in vitro*

Most research on LAMA2-CMD has been performed using *in vivo* models^11,12^ and even though this is of critical importance, the severity and complexity of the disease make difficult the identification of the precise disease mechanisms that underlie the onset of LAMA2-CMD. This is particularly relevant since the vast majority of previous studies used models where the secondary events of the disease were already present and therefore were likely to mask the primary events. To tackle this issue, we have generated an *in vitro* model for LAMA2-CMD using the myoblast cell line C2C12 (**Figure 1A**), which has been extensively used as an *in vitro* model to study myogenesis^4,20–22^. In this cell line, *Lama2* was deleted using CRISPR/Cas9 technology and different single cell clones were established. Deletion of *Lama2* was accessed by quantitative PCR (qPCR) for each single cell clone, as represented in **Figure 1B**, and, from these, at least two single cell clones were selected to perform each experiment to ensure that the phenotype observed was due to *Lama2* deletion and not associated with potential off target or clonal effects. The characterization of the mutations was performed by Sanger sequencing of the PCR products covering the gRNA-Cas9 target regions in *Lama2* exons 4 and 9 (**Supplementary Figures 1A-C**).

**Figure 1.**
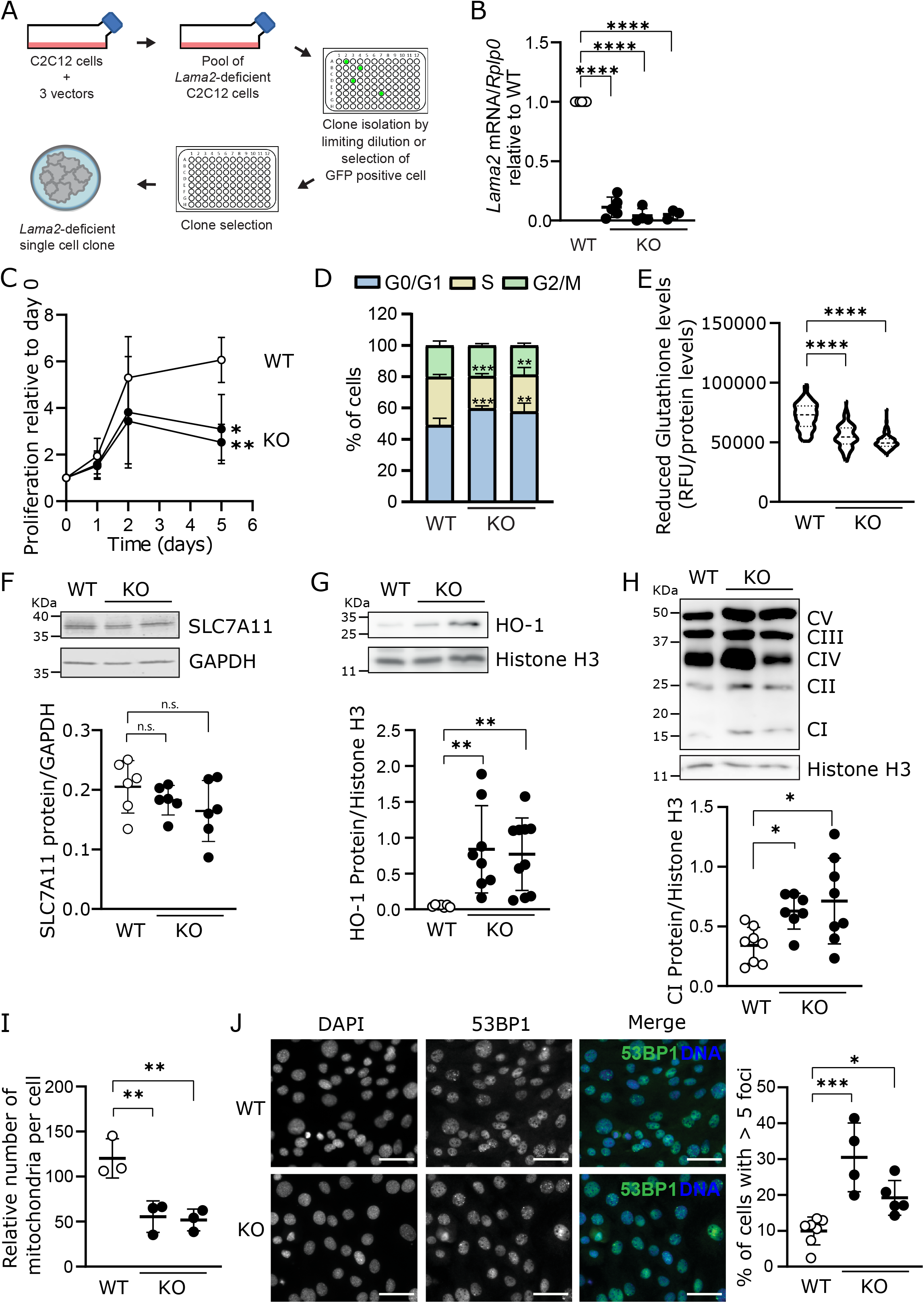
*Lama2*-deficiency impairs proliferation, possibly due to cell cycle arrest, and triggers DNA damage and oxidative stress. **A)** Schematic representation of *Lama2* knockout cell lines generation using CRISPR-Cas9 technology. **B)** C2C12 wildtype (WT) cells and three independent *Lama2* knockout single cell clones (KO) were harvested, the RNA was extracted and the expression of *Lama2* was analyzed by qPCR. *Lama2* expression levels were normalized to the housekeeping gene *Rplp0* and then for WT. n=3-7 independent samples per group collected independently. **C)** WT and KO C2C12 cells were plated, and cell proliferation was monitored on days 0, 1, 2 and 5 using Resazurin. Data was plotted relative to day 0, in order to analyze proliferation rate. n=3 independent experiments, each with 4 technical replicates. **D)** WT and KO C2C12 cells were stained with propidium iodide (PI) and analyzed by flow cytometry. The percentage of cells in G1/G0, S and G2/M cell cycle phase is represented. n=3-5 experiments per cell line. **E)** To measure the levels of reduced glutathione, WT and KO C2C12 cells were incubated with 40 µM of monochlorobimane for 30 minutes, and fluorescence was measured (Excitation: 390 nm; Emission: 490 nm). Fluorescence levels were normalized for protein levels (BCA quantification method). n=3-4 independent experiments, each with 36 independent measurements. **F-H)** WT and KO C2C12 cells were harvested for western blot analysis with anti-SLC7A11 **(F)**, anti-HO-1 **(G)**, or OXPHOS (CI -complex I; CII - complex II; CIII - complex III; CIV - complex IV; CV - complex V) **(H)** antibodies. Histone H3 or GAPDH was used as loading control. Two KO lanes represent experiments performed with two independent *Lama2* knockout single cell clones. n=6-10 samples per group collected independently. Densitometry analysis is shown under each western blot. **I)** To obtain the relative number of mitochondria per cell, WT and KO C2C12 cells were harvested, and total DNA was extracted and analyzed by qPCR. The ratio between *Nd1* DNA expression (encoded in the mitochondria) and *Hk2* DNA expression (encoded in the nucleus) was calculated and used as a proxy for the mitochondria number per cell. n=3 independent experiments. **J)** Representative images of WT and KO C2C12 cells fixed and processed for immunofluorescence with anti-53BP1 antibody, as DNA damage marker. Counterstaining of nuclei was performed with DAPI. n=4-7 independent experiments. Scale bar 50 µm. Quantification of the number of cells with more than five 53BP1 foci was performed. Images of two *Lama2* knockout single cell clones were analyzed separately. Statistical analysis was performed using Ordinary One-way ANOVA Dunnett’s multiple comparisons test for B), E) and H-J) and Two-way ANOVA with Tukey’s multiple comparisons test for C) and D). P-value: * p<0.05, ** p<0.01, *** p<0.001, **** p<0.0001. ROUT method was used to identify outliers.

Our previous data, using the *dy^W^* mouse model for LAMA2-CMD, showed that the disease onset was characterized by a significant reduction in the number of MuSCs and myoblasts^19^. To test if *Lama2*-deficiency affects cell proliferation, we compared *Lama2* knockout and wildtype C2C12 cells and observed a significant decrease in proliferation in the *Lama2* knockout cells (**Figure 1C**). This proliferation defect may be explained by an arrest of *Lama2* knockout cells in G1-phase of the cell cycle and a concomitant reduction in S-phase (**Figure 1D**, **Supplementary Figure 1D**).

### Absence of *Lama2* triggers oxidative stress and DNA damage

Induction of cell cycle arrest may be caused by different insults, for example oxidative stress and DNA damage^23^. Considering the important role of oxidative stress and mitochondrial dysfunction in LAMA2-CMD pathology^15–18,24^, we analyzed different markers for these insults in our *in vitro* model (**Figures 1E-H**). To assess the overall redox status of *Lama2*-deficient cells, we measured the levels of reduced glutathione (GSH), an important endogenous antioxidant, and found that they were significantly lower in *Lama2* knockout cells in comparison to their wildtype counterparts (**Figure 1E**), suggesting an increase in the oxidative status of *Lama2* knockout cells. The decrease in GSH was not likely associated with defective glutathione synthesis, as indicated by the levels of GCLC and GCLM (**Supplementary Figure 2A**), the two subunits of the enzyme catalyzing the rate-limiting step in the biosynthesis of GSH. However, we found that the cystine/glutamate transporter SLC7A11 was reduced, even though not significantly, in *Lama2*-deficient cells (**Figure 1F**). This possibly indicates an impairment in the transport of cystine, and consequently of the availability of cysteine, a precursor for glutathione synthesis. In accordance with the lower levels of GSH, we also found that the stress-induced heme oxygenase 1 (HO-1) was upregulated in *Lama2* knockout cells in comparison to their wildtype counterparts (**Figure 1G**).

Mitochondrial electron transport chain is an important source of reactive oxygen species (ROS) and mitochondrial dysfunction is therefore closely linked to oxidative stress^25^. To investigate if mitochondrial function was altered *in vitro*, we analyzed the levels of the oxidative phosphorylation (OXPHOS) complexes (**Figure 1H**, **Supplementary Figure 2B**). Complex I was increased in *Lama2*-deficient cells (**Figure 1H**), while the remaining complexes remained unchanged (**Supplementary Figure 2B**). Complex I is a key source of ROS production in the mitochondria, and increased levels of complex I have been associated with aging^26,27^. Importantly, decreased mitochondrial number has also been linked to skeletal muscle aging^28^. Therefore, we next analyzed the number of mitochondria in *Lama2*-proficient *vs*. deficient cells and found a significant reduction in the number of mitochondria in *Lama2*-deficient cells (**Figure 1I**). These data give further support to the notion that *Lama2*-deficient cells display increased ROS, which compromises mitochondrial integrity and may impair cell proliferation.

To further tackle the possible causes linked to defective proliferation and cell cycle arrest, we investigated whether the absence of *Lama2* could also lead to DNA damage, which can be triggered by different insults including increased ROS levels^13^. To evaluate DNA damage, we analyzed the percentage of cells displaying more than five 53BP1 foci and found that this percentage was significantly increased in *Lama2*-deficient cells when compared to their wildtype counterparts (**Figure 1J**). Hence, a significant increase in DNA damage may also be involved in the early stages of *Lama2*-deficiency.

### *Lama2*-deficiency disrupts normal cell fate mechanisms

Considering that our data suggests that *Lama2*-deficiency may lead to proliferation defects, increased oxidative stress and increased DNA damage in C2C12 cells (**Figure 1**), we went on to test if these changes could lead to the induction of apoptosis (**Supplementary Figure 2C**). Comparison of mRNA expression of the pro-apoptotic *Bax* gene and the anti-apoptotic *Bcl2* gene between *Lama2*-deficient and -proficient C2C12 cells did not show significant differences (**Supplementary Figure 2C**). Since previous studies have shown that autophagy is important to prevent MuSCs senescence and apoptosis^29,30^ but also for a correct myogenesis and muscle differentiation^31^, we tested if *Lama2-*deficiency could lead to impaired autophagy. For that we analyzed the levels of the autophagosome marker LC3 and found that LC3-II was significantly decreased in *Lama2*-deficient C2C12 cells in comparison to the wildtype (**Supplementary Figure 2D**). This reduction was also evident for LC3-I, which was frequently not detectable in *Lama2-*deficient cells. These findings suggest a defective autophagosome formation in *Lama2*-deficient cells.

### Differentiation is impaired in the *Lama2*-deficient cells

Since we observed a decrease in proliferation rate, we decided to analyze myoblast differentiation using this model. Generally, decreased proliferation or cell cycle arrest is linked to a change in the differentiation program, as these processes are sequential^32^. However, if a defective myoblast differentiation is observed, it would impair the formation, growth and/or the stability of muscle fibers. To analyze if differentiation is impaired in the absence of *Lama2*, we induced differentiation in *Lama2*-deficient and - proficient C2C12 cells *in vitro* and followed the formation of myotubes for 14 days (**Supplementary Figure 3A**). To avoid the confounding effect of the delayed proliferation observed in the *Lama2*-deficient cells (**Figure 1C and D**), the differentiation experiments were performed using normal proliferation medium (containing 10% FBS) and cells were allowed to differentiate by cell-cell contact. *Lama2*-proficient cells formed evident myotube-like structures by day 5, while similar structures could only be observed by day 14 in *Lama2*-deficient cells (**Supplementary Figure 3A**). To determine if the aligned cellular structures observed under the brightfield microscope were indeed myotubes, we stained *Lama2*-deficient and - proficient cells with an anti-myosin heavy chain (MyHC) antibody, a protein synthesized by myotubes, but not by myoblasts (**Figures 2A, B**). MyHC was detected in wildtype cells already at day 5 post-differentiation (**Figure 2A**), confirming the formation of myotubes, defined as cells with more than 2 nuclei. In contrast, *Lama2*-deficient cells presented rare and insipient myotube formation with only two nuclei at day 5 (**Figures 2A, A’**). Despite the increase in the number of aligned myoblasts on day 14, the number of fibers per total number of nuclei was still significantly reduced in the *Lama2*-deficient cells, and the vast majority of MyHC-positive cells had only one nucleus (**Figures 2B, B’**). This indicates a severe differentiation defect in the absence of *Lama2*. To better understand which mechanisms were implicated in this differentiation defect, we analyzed the expression of several myogenic markers in C2C12 cells grown under proliferation conditions (**Figure 3**). The mRNA expression of the MRF myogenin (*Myog*) (**Figure 3A**) was significantly lower in *Lama2*-deficient cells in comparison to their wildtype counterparts. Moreover, both the number of MYF5-positive nuclei (**Figure 3B**) and MYF5 nuclear protein levels (**Figure 3C**) were significantly reduced in *Lama2*-deficient cells compared to wildtype cells. This decrease was concomitant with an increase in NFIX nuclear protein levels (**Figure 3C**), and number of NFIX-positive nuclei (**Figure 3D**). NFIX is a transcription factor that is responsible for the transition between embryonic and fetal myogenesis^4,33^, and has been shown to be highly expressed in C2C12 cells^20^. These results suggest that increased levels of NFIX may cause a shift towards a faulty fetal myogenesis, compromising the skeletal muscle differentiation process. To determine whether later stages of myogenesis were affected by *Lama2*-deficiency, we analyzed the mRNA expression of the tubulin β6 chain (*Tubb6*) and myosin light chain (*Myl1*), both myogenin target genes, at day 5 post-differentiation (**Figures 3E, F**). Both genes, previously shown to be important players in differentiation^34–36^, were significantly reduced in *Lama2*-deficient cells when compared to wildtype cells (**Figures 3E, F**). Overall, these results suggest that *Lama2*-deficiency leads to a failure in promoting the necessary cytoskeletal organization, which is required for the transition from myoblast to myotubes.

**Figure 2.**
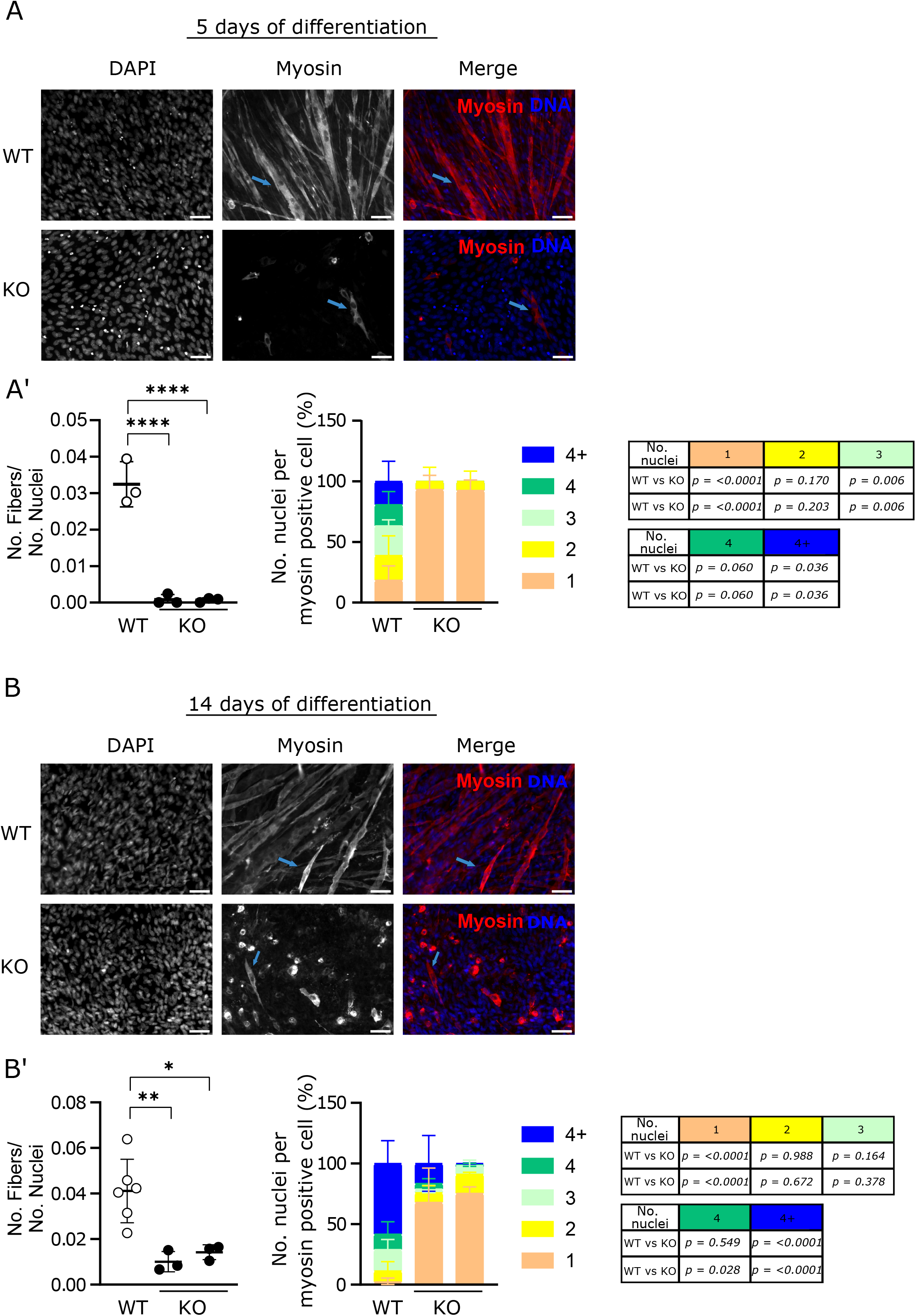
*Lama2* knockout cells show impaired differentiation. **A)** C2C12 wildtype (WT) and two independent *Lama2* knockout single cell clones (KO) were cultured for 5 days to induce differentiation and then fixed and processed for immunofluorescence with anti-myosin heavy chain antibody. Representative images are shown. Counterstaining of nuclei was performed with DAPI. n=3 independent experiments. Scale bar 50 µm. Full blue arrows indicate myosin positive cells. **A’)** The ratio between the number (No.) of fibers (myosin positive cells with 2 or more nuclei) per total number of nuclei was quantified (left panel), as well as the number of nuclei per myosin positive cell (middle panel) and respective p-values (right panel). **B)** C2C12 wildtype (WT) and two independent *Lama2* knockout single cell clones (KO) were cultured for 14 days in order to induce differentiation and then fixed and processed for immunofluorescence with anti-myosin heavy chain antibody. Representative images are shown. Counterstaining of nuclei was performed with DAPI. n=3-6 independent experiments. Scale bar 50 µm. Full blue arrows indicate myosin positive cells. **B’)** The ratio between the number (No.) of fibers (myosin positive cells with 2 or more nuclei) per total number of nuclei was quantified (left panel), as well as the number of nuclei per myosin positive cell (middle panel) and respective p-values (right panel). Statistical analysis in A’) and B’) was performed using Ordinary One-way ANOVA Dunnett’s multiple comparisons test. P-value: * p<0.05, ** p<0.01, *** p<0.001, **** p<0.0001.

**Figure 3.**
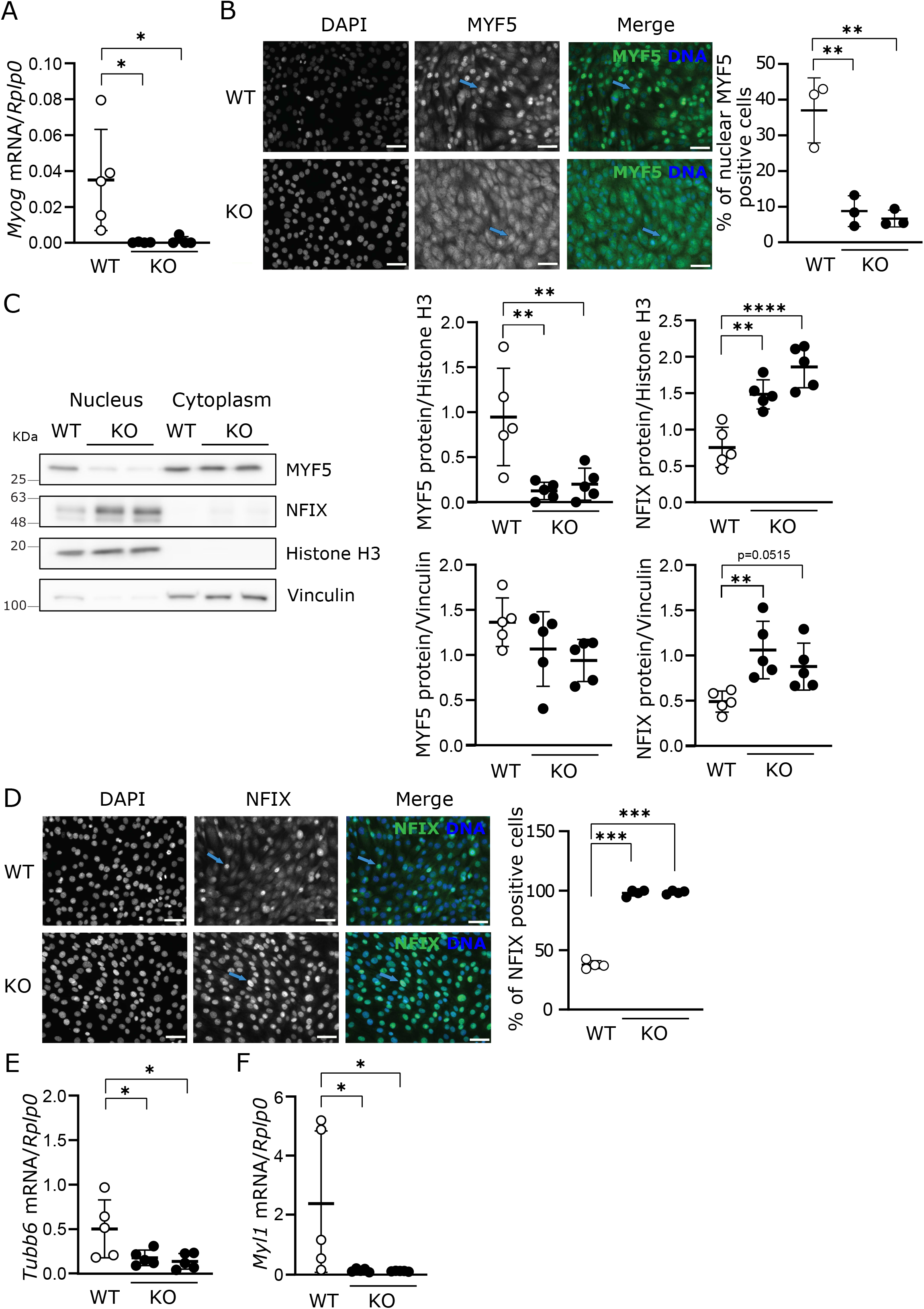
*Lama2* knockout cells show impaired differentiation and alterations in MYF5 and NFIX pathways. **A)** C2C12 wildtype (WT) cells and two independent *Lama2* knockout single cell clones (KO) were harvested, the RNA was extracted and the expression of *Myogenin* (*Myog*) was analyzed by qPCR. Expression levels were normalized to the housekeeping gene *Rplp0*. n=4-5 independent samples per group collected independently. **B)** Representative images of WT and KO C2C12 cells fixed and processed for immunofluorescence with anti-MYF5 antibody are shown. Counterstaining of nuclei was performed with DAPI. Images of two *Lama2* knockout single cell clones were analyzed separately. Quantification of the number of nuclear MYF5 positive cells is shown on the right. Blue arrows indicate MYF5 nuclear staining. Scale bar 50 µm. **C)** WT and KO C2C12 cells were harvested and the nuclear and cytoplasmatic fraction were separated by cell fractionation. Protein was extracted from each fraction and analyzed by western blot with anti-MYF5 and anti-NFIX antibodies (left panel). Histone H3 (nuclear phase) and vinculin (cytoplasmic phase) were used as loading controls. Two KO lanes represent experiments performed with two independent *Lama2* knockout single cell clones. n=5 samples per group collected independently. Densitometry analysis is shown on the right. **D)** Immunofluorescence analysis similar to the one described in B) using anti-NFIX antibody. Images of two *Lama2* knockout single cell clones were analyzed separately. n=3 independent experiments. Blue arrows indicate NFIX nuclear staining. Scale bar 50 µm. Quantification of the number of nuclear NFIX positive cells is shown on the right. **E)** Myogenin target genes *Tubb6* and *Myl1* were analyzed by qPCR in WT and KO C2C12 cells after 5 days in culture to induce differentiation. Expression levels were normalized using the housekeeping gene *Rplp0*. n=5 samples per group collected independently. Statistical analysis in A) to E) was performed using Ordinary One-way ANOVA by Dunnett’s multiple comparisons test. P-value: * p<0.05, ** p<0.01, *** p<0.001, **** p<0.0001.

### Absence of *Lama2* leads to a significant downregulation in gene expression

Considering our *in vitro* findings with the C2C12 LAMA2-CMD cellular model, highlighting the importance of *Lama2* in myoblast differentiation (**Figures 2,3 and Supplementary Figure 2**), we decided to perform an RNA sequencing (RNAseq) analysis on muscle fibers isolated from E17.5 fetuses, the developmental timepoint where the first signs of the disease have been reported^19^. The RNAseq analysis of muscle fibers revealed a profound downregulation of gene expression with 4958 genes downregulated in the *dy^W^* fetuses, when compared to the wildtype (adjusted p-value 0.01, Log2 fold change of ±1.5), while only 181 genes were found to be upregulated (**Figure 4A**). This indicates that lack of *Lama2* has a severe impact on the gene expression program of muscle fibers, which may affect their stability. To gain further insights into the pathways altered in the absence of *Lama2,* we performed a functional analysis. This revealed that downregulated genes were significantly associated with pathways linked to plasma membrane structure and function and transmembrane signaling, and also with intermediate filament organization and mechanisms involving maintenance of cellular redox status (**Figure 4B**). This supports the notion that absence of the α2 chain of laminin causes substantial structural changes in the link between basement membrane and cell cytoskeleton. On the other hand, genes that were upregulated were involved in protein and RNA processing and metabolism, as well as in collagen-containing extracellular matrix (**Figure 4C**). This increase in collagen is in accordance with previous findings in post-natal/adult muscle showing that fibrosis is a central hallmark of LAMA2-CMD pathogenesis^37,38^.

**Figure 4.**
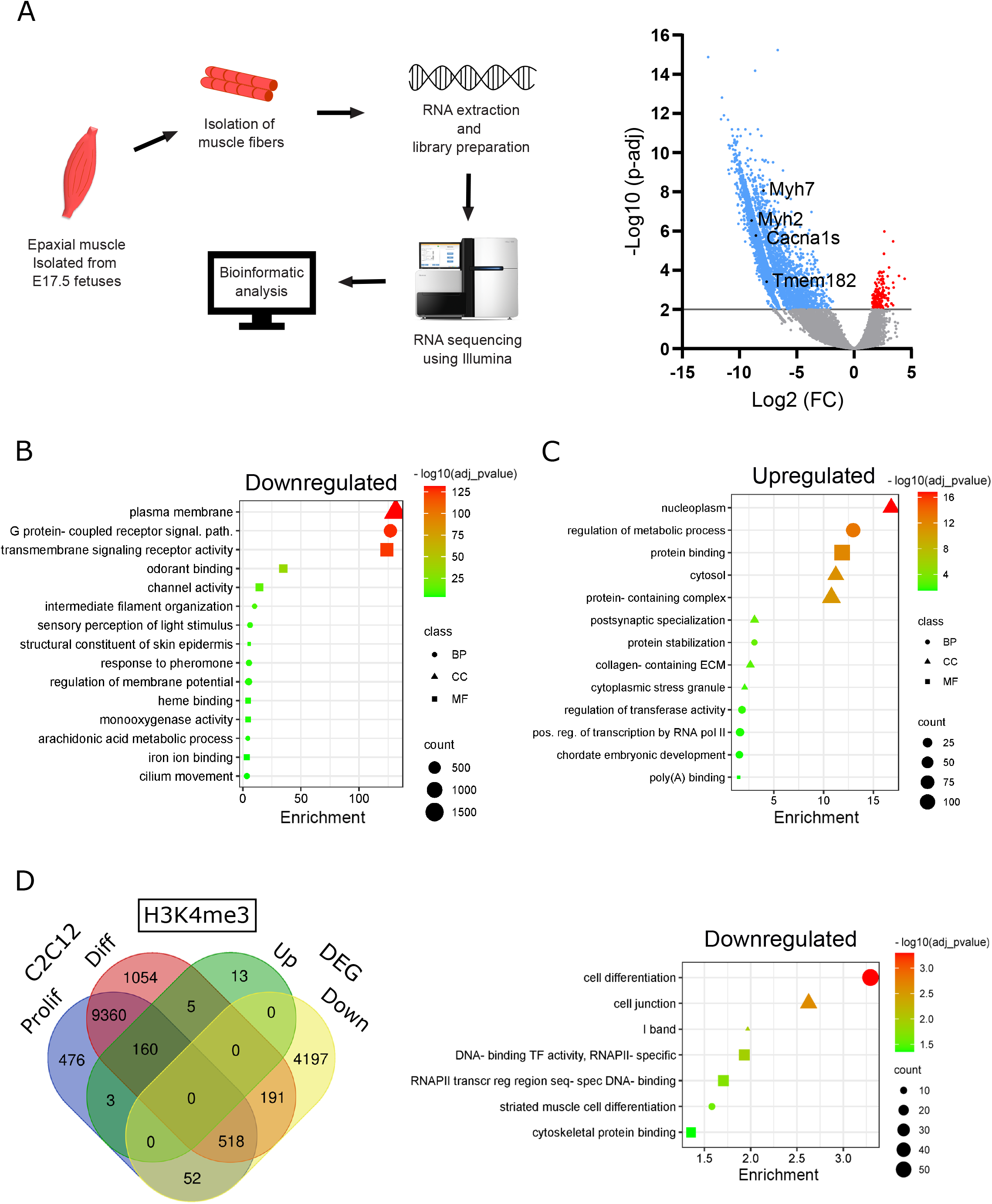
RNA sequencing analysis of muscle fibers from wildtype vs *dy^W^* fetuses at E17.5. **A)** Muscle fibers were extracted from wildtype (WT) and *dy^W^* fetuses at E17.5 and sent for RNA sequencing analysis. On the left is a schematic representation of the methodology used for the RNAseq experiments. On the right, the volcano plot of the genes differently expressed on the *dy^W^* fetuses when compared to WT is shown. In blue are represented the downregulated genes, in grey the non-significant and in red the upregulated. **B-C)** Functional analysis of genes downregulated (**B**) or upregulated (**C**) in the RNAseq analysis described in (A). Counts indicate the number of genes found in each category. Class indicates the functional analysis class BP (biological process, circles), CC (cellular component, triangle) and MF (molecular function, square). Color indicated the scale for the –log10 (adjusted p-value). **D)** Venn diagram comparing the differentially expressed genes (DEGs) (adjusted p-value 0.01, log2 fold change +/-1.5) (described in A) with previously published ChIP-Seq database (doi:10.17989/ENCSR000AHO, ENCODE), where C2C12 myoblasts and C2C12 myocytes (differentiated cells) were analyzed by histone ChIP-Seq for different histone modifications. The Venn diagram here represented compares specifically the histone H3 trimethylation in lysine 4 (H3K4me3) as a marker for active gene expression. **E)** Functional analysis of the 191 genes with active expression in C2C12 myotubes but downregulated in the RNAseq analysis of muscle fibers (described in D). Counts indicate the number of genes found in each category. Class indicates the functional analysis class BP (biological process, circles), CC (cellular component, triangle) and MF (molecular function, square). Color indicated the scale for the –log10 (adjusted p-value).

### *Lama2* is crucial for gene expression regulating muscle cell differentiation

Given the possible defect in muscle differentiation, we next compared the differentially expressed genes (DEGs) in fetal *dy^W^* muscle fibers versus wildtype controls, with a previously published ChIP-Seq database (doi:10.17989/ENCSR000AHO, ENCODE), where C2C12 myoblasts and C2C12 myocytes (*i.e.* differentiated cells) were analyzed by histone chromatin immunoprecipitation sequencing (ChIP-Seq) for different histone modifications. In particular, we compared the DEGs from our muscle fiber RNAseq with C2C12 myoblast and C2C12 myocyte ChIP-seq analysis for histone H3 trimethylation on lysine 4 (H3K4me3) (**Figure 4D**), a gene activation marker. One hundred and ninety-one genes were activated specifically in C2C12 myocytes (*i.e.* under differentiation conditions) and downregulated in *dy^W^* fetal muscle fibers (**Figure 4D, Supplementary Table 1**). Functional analysis showed that these genes are critical for cell differentiation, including striated muscle cell differentiation, as well as cytoskeletal structure (**Figure 4E**). These data strongly support the hypothesis that muscle fiber differentiation is also impaired in *dy^W^* fetuses.

To further expand our analysis on the pathways that were altered in *dy^W^ vs.* wildtype fetal muscle fibers, we compared the DEGs of wildtype vs. *dy^W^*muscle fibers with gene from different ontologies with relevance to our findings (**Figure 5, Supplementary Table 2-7**). This analysis showed perturbations in actin cytoskeleton organization and muscle cell differentiation, including the downregulation of *Cacna1s*, *Myh7* and *Tmem182* (**Figures 5A, Supplementary Table 2-3**), some of the top downregulated genes identified in our RNAseq analysis (**Figure 4A**). Additionally, we also compared DEGs with cell cycle and DNA repair gene ontologies, where genes such as the cell cycle regulator *E2f1* and the DNA repair genes *Exo1* and *Lig4* were found to be downregulated (**Figures 5B, Supplementary Table 4-5**). Comparison of DEGs with genes involved in oxidative stress response and mitochondrion organization revealed downregulation of different antioxidant genes such as *Gpx2*, *Gpx5*, *Gpx6* and *Nqo1* (**Figures 5C, Supplementary Table 6-7**). Collectively these data show that the absence of *Lama2* has a key impact on multiple pathways.

**Figure 5.**
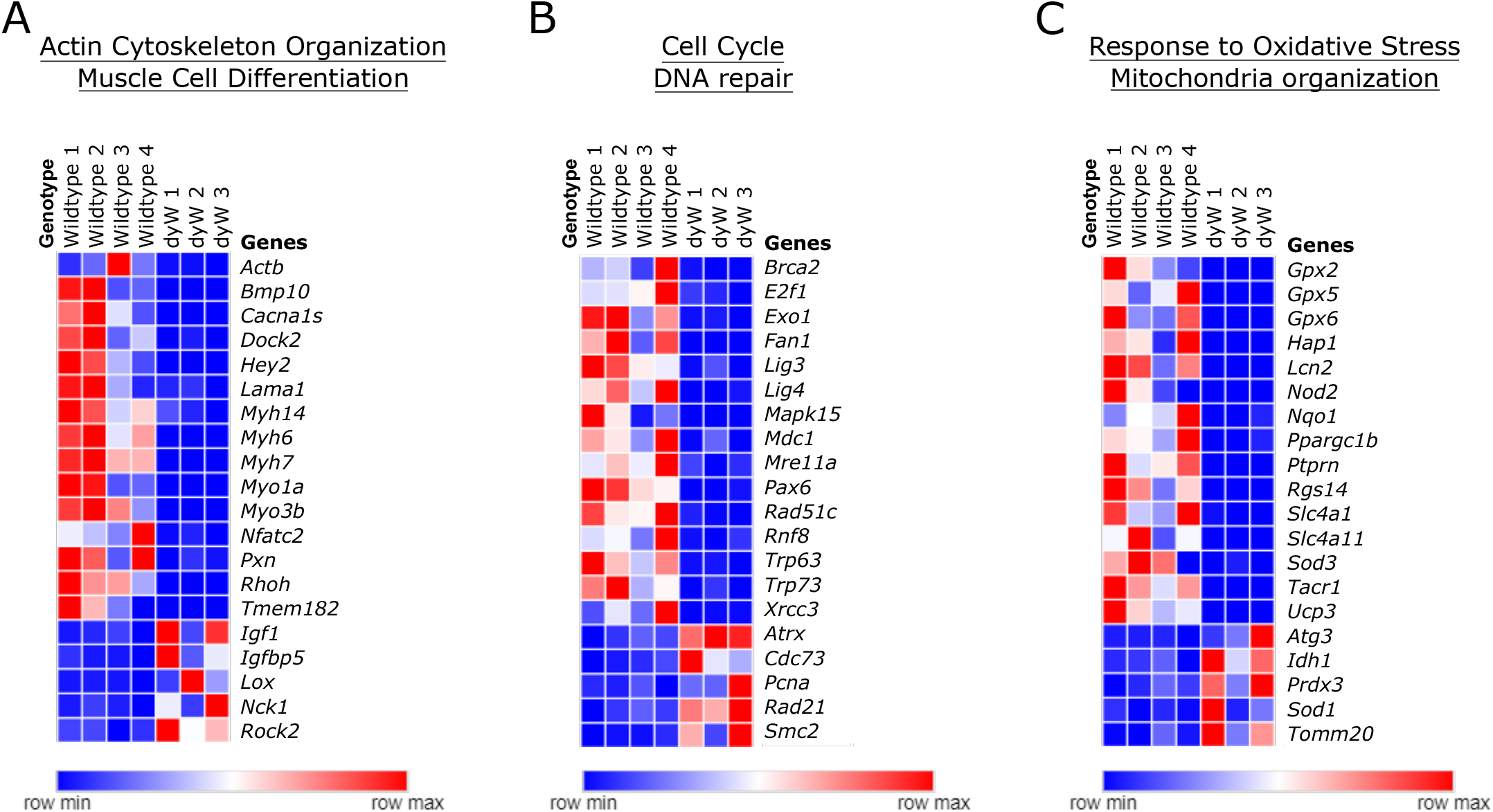
*Lama2*-deficiency alters the expression profile of different core cellular mechanisms. Heatmaps of 20 representative genes (15 downregulated and 5 upregulated) from the Venn diagram analysis comparing the differentially expressed genes (DEGs) (adjusted p-value 0.05, log2 fold change +/-1.5) of wildtype vs. *dy^W^* muscle fibers with the following gene ontologies: (A) GO:0030036 Actin Cytoskeleton Organization and GO:0042692 Muscle Cell Differentiation. (B) GO:0007049 Cell Cycle and GO:0006281 DNA repair. (C) GO:0006979 Response to Oxidative Stress and GO:0007005 Mitochondria organization.

### *Myh2*, *Myh7*, *Tmem182* and *Cacna1s* are downregulated in the absence of *Lama2*

To validate the important defect in muscle cell differentiation identified in our RNAseq analysis (**Figure 4**), we analyzed four top hits: the fast embryonic MyHC gene *Myh2*, previously shown to be implicated in skeletal muscle differentiation, and the embryonic slow myosin and myocyte marker (*Myh7*)^39–42^, the MYOD target gene *Tmem182*, a negative regulator of muscle growth^43^, but required for muscle cell fusion^44^, and the α1Ls subunit of voltage-gated L-type Ca^2+^ channel (Cav1.1) (*Cacna1s*), recently shown to be required for muscle fiber maturation^45^ (**Figure 6**). Comparison of RNAseq normalized counts (**Figures 6A-D**) with qPCR of isolated muscle fibers from E17.5 wildtype and *dy^W^* fetuses (**Figures 6E-H**) confirmed the significant downregulation of these genes, providing further evidence of a defective muscle differentiation and fusion signature in *dy^W^* fetuses. In addition, we analyzed whether the expression of these genes was also impaired in our *in vitro* model for LAMA2-CMD under differentiation conditions. After 5 days of differentiation, *Tmem182* and *Cacna1s* were significantly downregulated in *Lama2*-deficient cells (**Figures 6I-J**), while no significant difference was observed in *Myh7* expression, despite a tendency to be reduced in the absence of *Lama2* (**Supplementary Figure 3B**). *Myh2* expression did not show differences under our experimental conditions (**Supplementary Figure 3B**). However, *Myh7* was significantly decreased after 14 days of differentiation (**Figure 6K**), consistent with the defective differentiation phenotype found *in vitro* (**Figures 2,3 and Supplementary Figure 3**).

**Figure 6.**
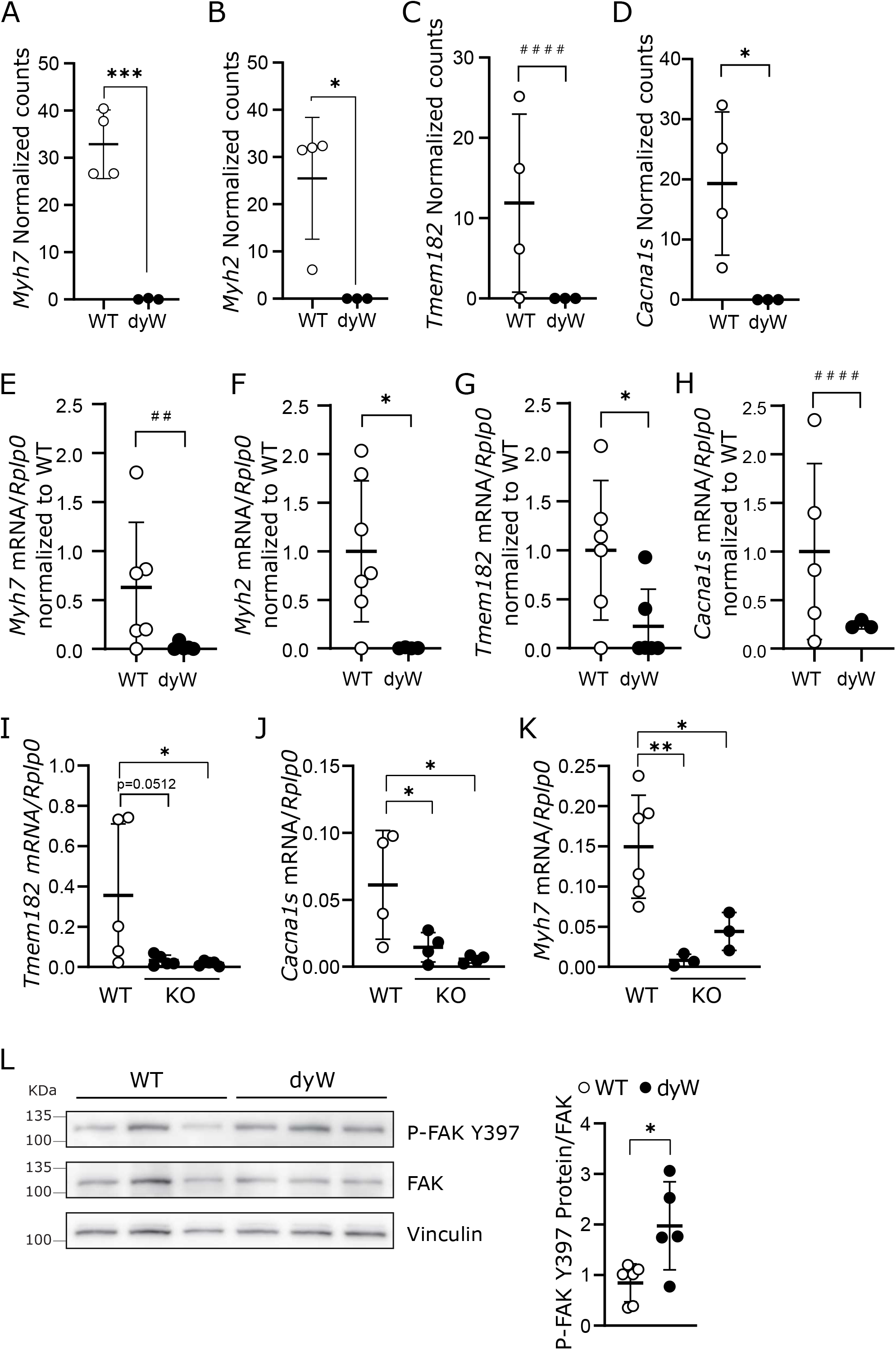
Expression of genes linked to cell differentiation. **A-D)** Normalized counts of genes selected from the analysis in Figure 4 related to cell differentiation (*Myh7* **(A)**, *Myh2* **(B)**, *Tmem182* **(C)**, and *Cacna1s* **(D)**). n=3-4 independent samples per genotype. **E-H)** To validate the data shown in A-D, epaxial muscles were collected from wildtype and *dy^W^*fetuses at E17.5 and muscle fibers were separated. RNA was extracted from muscle fibers and cDNA synthesized. The expression of genes selected from the analysis in Figure 4 were analyzed by qPCR (*Myh7* **(E)**, *Myh2* **(F)**, *Tmem182* **(G)**, *Cacna1s* **(H)**). n=3-7 independent samples per genotype. **I-K)** C2C12 wildtype (WT) cells and two independent *Lama2* knockout single cell clones (KO) were cultured for 5 **(I-J)** or 14 days **(K)** to differentiate and then harvested. RNA was extracted and the expression of *Tmem182* **(I)**, *Cacna1s* **(J)** and *Myh7* **(K)** was analyzed by qPCR. n=3-6 independent samples per group collected independently. **L)** Epaxial muscles were collected from wildtype and *dy^W^* fetuses at E17.5, the protein extracts were analyzed by western blot and the membranes were probed with anti-FAK Y397 and anti-FAK antibodies (left panel). Vinculin was used as the loading control. Densitometry analysis is shown on the right. n=5-6 fetuses per genotype. Statistical analysis was performed using Unpaired T-test (p-value * p<0.05; *** p<0.001)) or F test to compare variances (p-value ^##^ p<0.01; ^####^ p<0.0001) in A-H) and L) and with Ordinary One-way ANOVA in I-K). P-value: * p<0.05, ** p<0.01.

Since TMEM182 has been shown to negatively regulate muscle growth via its direct interaction with integrin β1 (ITGB1) and that its downregulation induced the activation of focal adhesion kinase (FAK) signaling^43^, we analyzed FAK phosphorylation (P-FAK) in whole muscle of E17.5 wildtype and *dy^W^* fetuses and found that P-FAK levels were higher in *dy^W^* (**Figure 6L**), consistent with *Tmem182* downregulation. At the same time, we also observed an increase in the expression in *Itgb1* in E17.5 *dy^W^* vs. wildtype in our RNAseq data (**Supplementary Figure 3C**). This is in line with previous reports^46^, indicating that lack of functional *Lama2* alters the stability of integrin β1, the β chain of the integrin α7β1, a central laminin 211 transmembrane receptor.

### Cell cycle defects, increased DNA damage and altered redox regulation are present at the onset of LAMA2-CMD

Considering that our data using the *in vitro* C2C12 model (**Figures 1**) and the RNAseq analysis of isolated E17.5 muscle fibers (**Figure 4, 5**) suggest that additional pathways other than cell differentiation and fusion are implicated in the onset of LAMA2-CMD, we validated cell proliferation, DNA damage, oxidative stress and mitochondria dysfunction in E17.5 whole muscles (**Figure 7**). mRNA expression analysis of the cell cycle inhibitors *Cdkn1a* (encoding p21) and *Cdkn1c* (encoding p57) showed a significant increase in fetal muscle of *dy^W^* in comparison to wildtype fetuses, consistent with defective cell cycle regulation (**Figure 7A**). Similarly, levels of DNA damage were also higher in fetal muscle of *dy^W^ vs.* wildtype fetuses, as determined by the increase in the phosphorylation of histone H2AX (γH2AX) (**Figure 7B**). Despite the significant downregulation observed for *Nqo1* gene expression in the RNAseq analysis (**Figure 5C, Supplementary Table 6**), the same was not observed when analyzing NQO1 levels in fetal muscle from *dy^W^* and wildtype mice at E17.5 (**Figure 7B**). Likewise, no changes were observed in the levels of the OXPHOS complexes (**Supplementary Figure 4A**). Considering that the onset of the disease was found to occur between E17.5 and E18.5^19^, we asked whether the defect in antioxidant response and mitochondria function could occur at E18.5. Our data showed that at E18.5 there was a significant decrease in the levels of NQO1 and an increase in the complex I of the mitochondria (**Figure 7C**), while the remaining OXPHOS complexes were not altered (**Supplementary Figure 4B**). These results are in accordance with our *in vitro* data (**Figure 1**), and further support the idea of an increase in oxidative stress at the onset of LAMA2-CMD.

**Figure 7.**
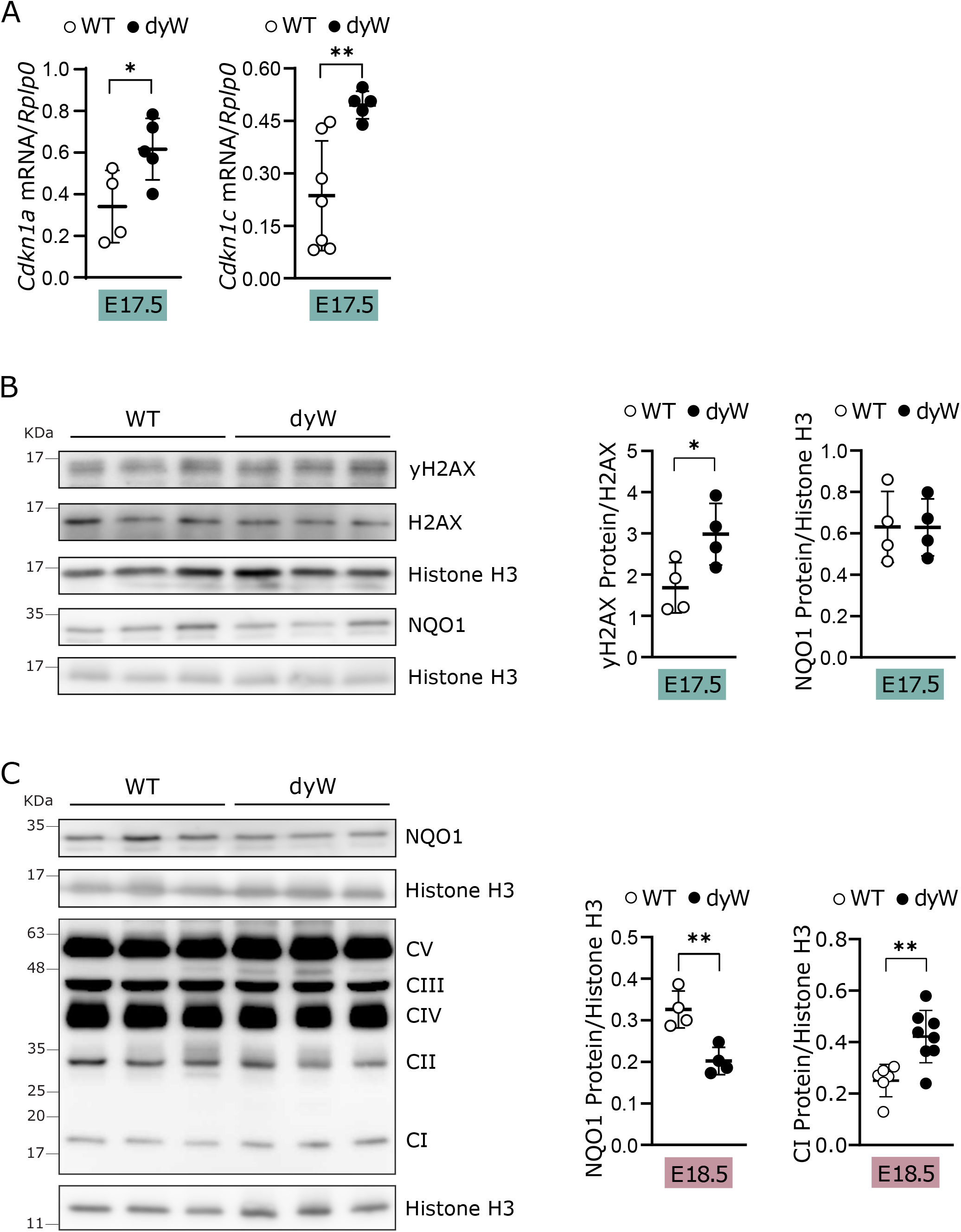
*Lama2*-deficiency leads to alteration in genes linked to cell cycle, triggers DNA damage and oxidative stress. **A)** Epaxial muscles were collected from wildtype (WT) and *dy^W^* fetuses at E17.5 and RNA was extracted. Expression of *Cdkn1a* and *Cdkn1c* was analyzed qPCR and normalized using the housekeeping gene *Rplp0*. n=4-7 fetuses per genotype. **B)** Epaxial muscles were collected from WT and *dy^W^* fetuses at E17.5, the protein extracts were analyzed by western blot, and the membranes were probed with anti-yH2AX, NQO1, and anti-H2AX antibodies (left panel). Histone H3 was used as the loading control. Densitometry analysis is shown on the right. n=4 fetuses per genotype. **C)** Epaxial muscles were collected from WT and *dy^W^* fetuses at E18.5, the protein extracts were analyzed by western blot and the membranes probed with NQO1 and OXPHOS (CI - complex I; CII - complex II; CIII - complex III; CIV - complex IV; CV - complex V) antibodies (left panel). Histone H3 was used as the loading control. Densitometry analysis is shown on the right. n=4-8 fetuses per genotype. Statistical analysis was done using Student’s t-test for A), B) and C). P-value: * p<0.05, ** p<0.01. ROUT method was used to identify outliers.

## DISCUSSION

Recent studies in mouse models of LAMA2-CMD by our laboratory and others, have shown that molecular and cellular alterations associated with this disease are detected early in life^14,19^. Moreover, an increase in Janus kinase/signal transducer and activator of transcription 3 (JAK-STAT3) signaling and a failure in normal muscle growth can be detected in fetal muscles of the *dy^W^* mouse model^19^, suggesting that the disease onset occurs between E17.5 and E18.5. Using both an *in vitro* model of LAMA2-CMD and the well-established *dy^W^* mouse model for this disease we have dissected the mechanisms underlying the onset of LAMA2-CMD and have found that a profound defect in cell differentiation and fusion, closely related to cytoskeletal changes, is at the core of LAMA2-CMD onset in these two models (**Figures 2-6 and Supplementary Figure 3**). The timing of the defect correlates with a stage at which MuSCs have just entered their niche under the basement membrane of myofibers, which in the case of *dy^W^* fetuses (and LAMA2-CMD patients) does not contain laminin 211. Several studies have already revealed the importance of laminin binding for cell proliferation and differentiation, in particular for stem cells^47,48^. Accordingly, MuSCs can display a proliferation defect in the absence of *Lama2*, as suggested by our *in vitro* data (**Figure 1C, D**), and by our previous findings showing a reduction in the number of PAX7^+^ cells in *dy^W^* fetuses^19^. Importantly, the absence of *Lama2* impairs cell differentiation and fusion *in vitro*, which is accompanied by a decrease in the myogenic regulatory factors *Myog* (**Figure 3A**) and MYF5 (**Figure 3B,C**), and an increase in the transcription factor NFIX (**Figure 3C,D**), which may reveal an unevenly entrance in the fetal myogeneisis^4,33^. Since NFIX expression should gradually increase during fetal development to assure a smooth transition from embryonic to fetal stages^49^, it is possible that this drastic alteration promotes cycles of regeneration and degeneration, exacerbating disease pathology, as previously reported in the context of α-sarcoglycan (*Sgca* null)- and dystrophin (*mdx)*-deficient dystrophic mice^50^. Therefore, while *in vitro* differentiation cannot occur in the absence of *Lama2* (**Figure 2 and Supplementary Figure 3A**), which had also been reported using *dy^W^* embryonic stem cells^51^, continuous myogenic program can generate muscle fibers *in vivo*. However, our RNAseq analysis showed that these fibers are severely compromised, with a profound downregulation of gene expression (**Figure 4A**). These results raise the possibility that treatment with histone deacetylase inhibitors, which promote gene expression, could have a positive effect on LAMA2-CMD, as previously shown in the context of other muscular dystrophies such as Duchenne muscular dystrophy^52,53^.

Comparison of the top downregulated and upregulated genes with a previously published H3K4me3 ChIP-Seq database from C2C12 myoblasts and C2C12 myocytes, allowed us to identify 191 genes that were expected to be active under differentiation conditions in C2C12 cells and were downregulated in our RNAseq analysis of E17.5 *dy^W^* muscle fibers (**Figures 4D, Supplementary Table 1**). These genes were involved in cell differentiation, including striated muscle cell differentiation, as well as cytoskeleton structure (**Figure 4E**). This further supports the idea of a differentiation and fusion defect. Among these genes, two were identified as top hits in our RNAseq analysis, *Tmem182* and *Cacna1s* (**Figure 4A**), and these findings were validated both *in vivo* and *in vitro* (**Figure 6**). Additionally, two key myosin heavy chain genes involved in muscle cell differentiation, *Myh7* and *Myh2,* were identified as being severely downregulated (**Figure 4A**). MYH7 is a slow myosin that plays a critical role during embryonic and fetal muscle development. As the development continues, other myosin chains start to be expressed, including the fast myosin MYH2, and are present from late fetal or early postnatal stages onwards^54^. Defective expression of *Myh7* and *Myh2* may compromise the formation and stability of *Lama2*-deficient myofibers (**Figures 2 and 6**). Moreover, this abnormal expression of myosin heavy chains may be related with the increased NFIX levels (**Figure 3C, D**), which is in accordance with previous studies showing that NFIX cooperates with SOX6 to repress the expression of MyHC-I (encoded by *Myh7*) during fetal muscle development^49^.

TMEM182 is a transmembrane protein with high expression in muscle and adipose tissues^55^. Previous studies revealed that TMEM182 plays a role in myogenesis^43,55^, where it negatively regulates muscle growth via its direct interaction with integrin β1^43^, while at the same time it is required for muscle cell fusion^44^. The absence of *Lama2* leads to *Tmem182* downregulation (**Figures 4A and 6**) and the consequent reduction in TMEM181-ITGB1 interaction is perhaps linked with the observed increase in P-FAK levels (**Figure 6L**). Even though FAK activation promotes differentiation and fusion^56^, the profound alterations observed in the absence of *Lama2* (**Figure 4A**) are in line with the generation of aberrant myotubes. One such alteration may be associated with the regulatory mechanisms that maintain intracellular calcium levels, as suggested by the significant decrease in *Cacna1s* (**Figures 4A and 6**), which has been studied in the context of muscle differentiation in C2C12 cells^57,58^ and has been associated with LAMA2-CMD disease pathology^17^. Moreover, the increase in mRNA expression of *Cacna1s* and other calcium-related genes coincided with that of myogenic factors such as *Myf5*, *Myod* and *Myog* ^57^. This is consistent with our findings showing that *Lama2*-deficiency leads to a simultaneous decrease in *Myog* (**Figure 3A**), MYF5 (**Figure 3B,C**) and *Cacna1s* (**Figure 6J**). Our *in vivo* findings from the RNAseq analysis (**Figure 4A**), and subsequent validation (**Figure 6H**), support the idea that *Cacna1s* expression is not only required for muscle differentiation, but also for its maturation^45^. Moreover, downregulation of *Cacna1s* has also been shown to occur in C2C12 cells treated with fibroblast growth factor 9 (FGF9)^58^, and correlated with defective differentiation. In contrast to our findings, this study shows that FGF9 leads to a shift towards a more proliferative phenotype^58^. In our study, is likely that the absence of *Lama2* produces much more drastic changes. In keeping with this notion, we have identified that defective proliferation, DNA damage and oxidative stress were also hallmarks of the disease onset, which were observed both in *in vitro* (**Figure 1**) and *in vivo* (**Figure 5 and 7**). Moreover, our *in vitro* data (**Supplementary Figure 2C**) and previous *in vivo* findings^19^ suggest that, even though apoptosis is a hallmark of LAMA2-CMD and its inhibition improved the outcome of the disease^59^, it seems to be a consequence of the disease rather than a primary cause. This result is consistent with the absence of apoptotic fibers observed in the *dy^3K^/dy^3K^* mouse model of LAMA2-CMD at E18.5^14^.

DNA damage and oxidative stress could also play a role in modulating muscle differentiation, since it has been shown that both DNA damage^60^ and increased expression of HO-1^61^ can inhibit C2C12 differentiation, the former by inhibiting muscle-specific genes^60^ and the latter by targeting striated muscle specific miRNAs, called myomiRs^61^. Additionally, the decreased levels of reduced glutathione observed in our *in vitro* model (**Figure 1E**) are consistent with the previously described aged MuSCs^62^, which may explain the exhaustion of the pool of MuSCs previously observed in *dy^W^* fetuses^19^. Moreover, our data suggests *Lama2*-deficiency triggers DNA damage, in both the *in vitro* model (**Figure 1J**) and *in vivo* in E17.5 fetuses (**Figures 5D and 7B**). This occurs before the observed changes in oxidative stress and mitochondrial composition, which occur at E18.5 (**Figure 7C**). Therefore, it is unlikely that the increase in DNA damage is a consequence of increased ROS levels, but rather an independent mechanism that can have an important contribution to the disease. Accordingly, DNA repair is decreased in differentiated myoblast/myotubes, in particular base excision repair, a DNA repair mechanism that is the first line of response against DNA oxidation^63^. In the absence of *Lama2,* this might be exacerbated by the downregulation of several DNA repair genes, including key genes involved in homologous recombination (e.g. *Brca2*, *Rad51c, Mdc1, Exo1*, *Mre11*) (**Figure 5B, Supplementary Table 5**), a central pathway in DNA double strand break repair. This downregulation of DNA repair genes and the differentiation-related decrease in base excision repair is likely to promote the increase in DNA damage that is further promoted by the accumulation of ROS contributing to myofiber degeneration. Additionally, the link between defective DNA repair/cell cycle and oxidative stress-associated mitochondrial dysfunction may also be potentiated by the increased expression of *Cdkn1a* in the absence of *Lama2* (**Figure 7A**). *Cdkn1a* encodes the key cell cycle regulator p21 and increased levels of p21 lead to mitochondria dysfunction, DNA damage and skeletal muscle dysfunction^64^. Possible mechanisms associated with the increase in oxidative stress may be the increase in mitochondrial complex I (**Figure 1H and Figure 7C**) and changes in intracellular calcium, since complex I has also been implicated in the regulation of calcium transport^65,66^. *In vivo*, an increase in complex I and reduction in the levels of the antioxidant protein NQO1 is not evident at E17.5, but rather at E18.5 (**Figures 7C,B**), suggesting that it may be a consequence of an initial dysfunction. This reinforces the idea that treatment with antioxidants is a potential therapeutic approach in the context of LAMA2-CMD, as proposed by other studies in post-natal stages^16^.

The overall dramatic impact caused by the absence of *Lama2* leads to profound alterations in important signaling pathways that are involved in the communication between intra- and extracellular environments, such as FAK (**Figure 6L**) and JAK/STAT3^19^. Another possibility is that faulty laminin 211-integrin binding alters cellular biomechanical properties. Several studies have shown that changes in ECM structure and rigidity (*i.e.* ECM stiffness), that can arise from the collagen-enriched ECM from *dy^W^* fetuses (**Figure 4C**), can impair proliferation^67^, impact nuclear stability and trigger DNA damage^68,69^ and lead to changes in mitochondrial function and fusion^70–72^. The changes in ECM composition are likely the first signs of fibrosis, which characterizes later stages of LAMA2-CMD^11,12^, but not its onset^19^.

In the present study, we provide the first evidence of the molecular and cellular processes underlying the onset of LAMA2-CMD in mouse models of the disease. Even though previous studies have reported proliferation and differentiation defects, as well as increased oxidative stress and metabolic alterations^11–19^, as important hallmarks for LAMA2-CMD disease pathology, we have for the first time linked some of these changes to disease onset during fetal development. Our results provide an important framework for studies that aim at targeting therapeutically the primary defect of this disease, which can hopefully increase the lifespan and quality of LAMA2-CMD patients. Additionally, since a significant number of alterations seem to be common between muscular dystrophies, these findings can also give important clues to the study of other less understood muscular dystrophies.

## ACKNOWLEDGEMENTS

The authors would like to acknowledge the core facilities of the Faculty of Sciences of University of Lisbon (FCUL) and Centre for Ecology, Evolution and Environmental Changes (cE3c) & CHANGE. SGM was supported by Fundação para a Ciência e Tecnologia (FCT, Portugal; 2022.10813.BD) and the COST (European Cooperation in Science and Technology) Action CA20121 (Bench to bedside transition for pharmacological regulation of NRF2 in noncommunicable diseases, BenBedPhar) (E-COST-GRANT-CA20121-a076558c), ARC by Fundação para a Ciência e Tecnologia (FCT, Portugal; CEECIND/01589/2017) and L’Oréal Portugal Medals of Honor for Women in Science 2019, and ST by Association Française contre les Myopathies (AFM) Téléthon (contract no. 23049). SGM, VR, CM, CPV, IF, BS, MP, DRF, PGS, GR, RZ, ARC and ST were supported by the donor Henrique Meirelles who chose to support the MATRIHEALTH Project (CC1036), Fundação para a Ciência e a Tecnologia Project (doi:10.54499/PTDC/BTM-ORG/1383/2020) and Unit Funding (doi:10.54499/UIDB/00329/2020). FH and FM were supported by centre grants to BioISI (doi:10.54499/UIDB/04046/2020) and individual grants (doi:10.54499/PTDC/FIS-MAC/2741/2021 and SFRH/BD/133220/2017, respectively) through Fundação para a Ciência e Tecnologia. Immunofluorescence experiments were supported by Microscopy Facility of Faculty of Sciences of the University of Lisbon (PPBI-POCI-01-0145-FEDER-022122). Mouse bred and maintenance were supported by CONGENTO LISBOA-01-0145-FEDER-022170 (FCT, Lisboa2020, Por2020, ERDF) and heterozygous crossing by FCUL animal house. Cell sorting of C2C12 cell line was performed at Flow Cytometry Facility of *Instituto de Medicina Molecular João Lobo Antunes* (IMM, Portugal), to whom we would also like to the technical support. Flow Cytometry analysis of C2C12 cell lines was performed at Margarida Gama-Carvalho Laboratory at FCUL. RNA sequencing was performed at the Genomics Unit of *Instituto Gulbenkian de Ciência* (IGC, Portugal), to whom we would like to thank for the technical support.

## AUTHOR CONTRIBUTION

A.R.C. wrote the manuscript with the contribution of S.G.M., V.F., D.R.F., and S.T., and A.R.C. and S.T. conceived the research. S.G.M., designed and carried out most of the experimental work. C.M., C.P.C., B.S. and D.R.F. generated the *in vitro* model. S.G.M., S.D.N, A.T.D.K. and I.F. performed the experiments and analysis related with oxidative stress. S.G.M., I.F. and M.P. performed the mitochondrial experiments and analysis. C.M., A.R.C. and S.G.M. performed the experiments and analysis related with cell cycle and proliferation. S.G.M., V.R. and C.P.C. performed the experiments and analysis related with differentiation. F.M. and F.H. performed the autophagy experiments and analysis. S.G.M., V.R., P.G.S., R.Z., A.R.C. and M.P. collected the samples sent to RNAseq and performed the RNAseq data validation. D.A. and A.R.C. performed the bioinformatic analysis of RNAseq data. P.G.S., A.R.C., V.R. and S.G.M. bred and maintained mice and collected fetal samples. All the authors contributed to the article with active discussion and revision and approved the final submission. All authors have read and agreed to the published version of the manuscript.

## DECLARATIONS OF INTERESTS

The authors declare that the research was conducted in the absence of any commercial or financial relationships that could be construed as a potential conflict of interest.

## MATERIALS AND METHODS

### Animals

Mice were bred and maintained under specific pathogen-free (SPF) conditions at the Instituto Gulbenkian de Cielncia (IGC) and heterozygous crossings were performed at FCUL (Ofício n° 99 0420/000/0009/11/2009). All experimental protocols were approved by the Ethics Committee of IGC, the Organization Responsible for animals’ welfare (Órgão Responsável pelo Bem*-estar dos Animais* (ORBEA)) at IGC and at FCUL and the Portuguese National Entity (Direcc□ão Geral de Alimentac□ão e Veterinária). Experimental procedures were performed according to the Portuguese (Portaria n° 1005/92, Decreto-Lei n° 113/2013 and Decreto-lei no1/2019) and European (Directive 2010/63/EU) legislations concerning housing, husbandry, and animal welfare. Heterozygous *dy^W^* C57BL/6 mice (gift from Eva Engvall via Paul Martin; The Ohio State University, Columbus, OH, USA) were crossed to obtain homozygous *dy^W^* mutants and wildtype fetuses^73^. Pregnant females were anesthetized with isoflurane, sacrificed by cervical dislocation and the uterine horns were removed and placed in ice-chilled PBS. Fetuses were removed from the uterine horns, maintained in ice-chilled PBS and decapitated. Tails from fetuses were collected for genotyping. For tail lysis, 25 mM NaOH / 0,2 mM EDTA was added to the samples and placed in thermocycler at 95°C for 30 min, then 40 mM Tris HCl pH 5,5 was added to the mixture in a 1:1 ratio and centrifuged for 3 min. For PCR, 1 μL of undiluted mixture was used per reaction. All PCR reactions were performed using Xpert Fast Hotstart DNA Polymerase according to the manufacturer’s instructions, under the following conditions: 94°C/2 min, 10 cycles/94°C/20 sec, 65°C/15 sec (−0.5°C/cycle), 68°C/10 sec, 28 cycles/94°C/15 sec, 60°C/15 sec, 72°C/10 sec. A single mix was prepared with the primers listed in the Key Resources Table (*Oligonucleotides – DNA genotyping*). Epaxial muscles were isolated from E17.5 and E18.5 wildtype and *dy^W^* fetuses and used in the different analysis or kept at −80°C for long-term preservation.

### Cell lines

C2C12 myoblast cell lines were grown in Dulbecco’s Modified Eagle’s Medium (DMEM), supplemented with 10% fetal bovine serum (FBS) and 1% of an antibiotic mixture: Penicillin (10000U/mL) and Streptomycin (10mg/mL) (complete or proliferation medium). The cell lines were maintained in culture at 37°C, 5% CO_2_ and constant humidity. When 70% of confluency was reached, trypsin-EDTA or TrypLE™ Express Enzyme was used to split the cells.

### Generation of *Lama2* KO C2C12 cell lines

The *in vitro* model for LAMA2-CMD was established through the deletion of *Lama2* gene from C2C12 cells by CRISPR-Cas9 (knockout C2C12 cell lines) using two gRNA targeting exons 4 and 9 of *Lama2* cloned in pRP[CRISPR]-Puro-hCas9-U6) (Key Resources Table *Oligonucleotides – gRNA oligos and cloning oligos*). Transfection of C2C12 cells was performed using Lipofectamine 3000 transfection reagent, according to manufacturer’s instructions. Approximately 48h after transfection, selection was performed using 3µg/mL puromycin for 48h. Upon selection single cell clones were isolated by limiting dilution (8 cells/mL). Alternatively, C2C12 cells were co-transfected with the plasmids carrying the gRNA and a GFP-expressing plasmid. In this second approach, GFP-positive cells were individually isolated through FACS (FACSAria III (BD Biosciences)). In both approaches, isolated clones were expanded in complete medium with 20% FBS. qPCR was performed to evaluate the *Lama2* deletion in C2C12 single cell clones (Key Resources Table *Oligonucleotides – qPCR oligos*), and Sanger sequencing was performed to confirm the gene editing (STABVIDA). Additionally, to confirm the presence of deletions, a PCR reaction was performed (see Key Resources Table *Oligonucleotides – gRNA oligos and cloning oligos* for primers and animal section on methods for PCR protocol). Transfection and clonal selection were repeated across time, considering the features found to be associated with *Lama2* deletion.

### Differentiation assay

Wildtype and *Lama2* knockout C2C12 cells were plated in DMEM complete medium at a density of 50 000 cells per well in a 24-well plate with coverslips and left to differentiate until day 5 or 14. Images were acquired on days 0, 2, 5, 7, 12 and 14 using an Optika IM-3LD4 with LED fluorescence under 20x objective lens coupled with a Canon EOS M200 camera. On day 5 or day 14, cells were fixed for immunofluorescence assay or collected for RNA extraction.

### Proliferation Assay

To analyze the proliferation of the different C2C12 cell lines, a resazurin assay was performed. For that, on the day before the experiment, wildtype and *Lama2* knockout C2C12 cells were plated in a 96-well plate (5000 cells per well). On the following day (day 0), cells were incubated in 1x Resazurin solution in complete DMEM medium for 1 h 20 min. Fluorescence (excitation filter 531/40 nm; emission filter 595 nm) was measured using Victor 3V plate reader (PerkinElmer) on days 0, 1, 2, and 5. The fluorescence levels were normalized to control and to day 0 for each cell line.

### Cell cycle analysis

To evaluate the cell cycle of the different C2C12 cell lines, cells and growth media were collected and washed with 1x PBS. Then resuspended 300 µL of PBS and after 700 µL of ice-cold absolute ethanol were added dropwise. Cells were fixed at −20°C for at least 24h. Immediately before the analysis by flow cytometry, cells were centrifuged, the ethanol was discarded, and the cells resuspended in 1x PBS with 5 mg/mL RNase for 5 min at room temperature (RT). After incubation, the cells were centrifuged and resuspended in 1x PBS with 50 μg/mL of Propidium iodide (PI). Flow cytometry analysis was performed using the BD FACSCalibur™ Flow Cytometer (BD Biosciences).

### Glutathione assay

Wildtype and *Lama2* knockout C2C12 cells were plated in a 24-well plate with a confluency of 70-90%. To measure the levels of reduced glutathione, cells were washed with serum free DMEM medium without phenol-red and then with Fluorobrite DMEM medium. After the washes, cells were incubated with 40 µM of Monochlorobimane diluted in Fluorobrite DMEM medium for 30 min. Fluorescence was measured using a plate reader (TECAN, Spark 10M) with an excitation wavelength of 390 nm and emission wavelength of 490 nm. For each well, 36 measurements of different sections of the well were obtained and then normalized for the protein levels of each well. For protein quantification, cells were recovered and lysed in lysis buffer (50 mM Tris pH6.8, 2% SDS and 10% Glycerol) and then protein levels were measured using the BCA quantification method.

### Fractionation assay

To separate the nuclear and cytoplasmatic fractions, a cell fractionation protocol was used^74^. Wildtype and *Lama2* knockout C2C12 cells were collected and washed in PBS and then resuspended in ice-cold fractionation buffer 10 mM HEPES pH 7.9, 10 mM KCl, 1.5 mM MgCl_2_, 0.34 M sucrose, 10% glycerol, 0,075% triton, 1mM DTT and protease inhibitor cocktail) and incubated on ice for 5 min. The samples were centrifuged 5 min at 1300 g at 4°C. The supernatant containing the cytoplasmic proteins was cleared by centrifugation for 20 min at 20 000 g at 4°C and then resuspended in 2x SDS-PAGE sample buffer (20% Glycerol, 4% SDS 100mM, Tris pH 6.8, 0.2% Bromophenol blue, and 100mM DTT) in a 1:1 ratio. The pellet containing the nuclear proteins was washed twice with ice-cold fractionation buffer, centrifuged 5 min at 1300 g at 4°C and resuspended in 2x SDS-PAGE sample buffer. Protein concentration was measured with Nanodrop1000 at 280 nm.

### RNA extraction and qPCR

Epaxial muscles collected from E17.5 and E18.5 fetuses or C2C12 cells were lysed using the tripleXtractor reagent and a tissue homogenizer (Retsch MM400 Tissue Lyser). The RNA was extracted according to the manufacturer’s instructions. RNA concentrations and quality were determined with Nanodrop1000. The Xpert cDNA Synthesis Kit was used to synthesize complementary DNA (cDNA) according to the manufacturer’s instructions. The cDNA obtained was stored at −20°C until further analysis by qPCR. qPCR reaction was performed with iTaqTM Universal SYBR® Green Supermix or Xpert Fast SYBR (Uni) Blue, according to manufacturer’s instructions, using CFX96TM Real-Time PCR Detection System (Bio-Rad). Transcript levels were normalized against *Rplp0* expression, and the fold change was calculated using ΔΔCt method. Primers used are listed in Key Resources Table *Oligonucleotides – qPCR oligos*.

### Quantification of the number of mitochondria per cell

Wildtype and *Lama2* knockout C2C12 cell lines were harvested, and total DNA was extracted using the protocol previously described by Quiros and colleagues^75^.

### Western Blot

To extract protein from muscle and C2C12 cells, samples were collected and resuspended in 2x SDS-PAGE sample buffer. Then, cells and tissues were homogenized with the tissue homogenizer (Retsch MM400 Tissue Lyser), when required further incubated with benzonase for 20 min, heated at 50°C for 10 min and centrifuged at maximum speed for 5 min. Protein quantification was measured with Nanodrop1000 at 280 nm. Protein extracts were separated with 10, 12 or 15% SDS polyacrylamide gel electrophoresis in running buffer (3,02 g Tris base, 14,42 g Glycine, 1 g SDS in 1 L distilled water) using the Mini-PROTEAN® Tetra electrophoresis system (Bio-Rad). Then, proteins were transferred to PVDF membranes on Mini Trans-Blot® Cell (Bio-Rad) with chilled Transfer Buffer (5.82g Tris, 2,93 g glycine in 1 L distilled water) and blocked with 5% milk in TBST (20 mM Tris, 150 mM NaCl, 0.1% Tween20 and distilled water, pH 7.4-7.6), with agitation. Protein loading was verified by GelCodeTM Blue Safe Protein Stain. Primary antibodies diluted in 2% bovine serum albumin (BSA) in TBST and 0.02% Sodium Azide were incubated overnight at 4°C with agitation (antibodies and dilutions used are listed in Key Resources Table). Secondary antibodies coupled with horseradish peroxidase (HRP) diluted in 5% powdered milk in TBST were incubated for 1h at RT, after being previously washed in TBST. Signal detection was obtained with Supersignal^TM^ West Pico Chemiluminescent Substrate HRP and images were acquired using the Amersham Imager 680 RGB (GE Healthcare). Quantifications were performed in Fiji software.

### Immunofluorescence

Wildtype and *Lama2* knockout C2C12 cell lines were plated in a 24-well plate with coverslips (50 000 cells per well). Cells were washed with 1x PBS and fixed for 10 min with 4% paraformaldehyde in PBS, at RT. After washing with PBS, cells were permeabilized in 0,1% Triton-X100 in PBS for 5 min and washed with PBS. Then, cells were blocked with blocking solution (1% BSA, 1% goat serum, 0.05% Triton-X100 in PBS) for 1h at RT and incubated with the primary antibody (see table S4) diluted in blocking solution overnight at 4°C. The following day, cells were washed with blocking solution and incubated with the secondary antibody (see Key Resources Table) diluted in blocking solution for 1h at RT. Nuclei were stained with DAPI (4′,6-diamidino-2-phenylindole) for 30 sec. Samples were mounted using Mowiol-DABCO mounting medium. Images were acquired using an Olympus BX60 fluorescence microscope under 20x objective lens coupled with Hamamatsu Orca R2 cooled monochromatic CCD camera.

### Fibers RNA sequencing and analysis

Epaxial muscle from wildtype and *dy^W^* fetuses at E17.5 were collected and digested in 37°C digestion buffer (dispase II 6U/mL and collagenase A 1U/mL in DMEM complete medium) with strong agitation for 45 min at 37°C. After incubation, the digestion buffer was inactivated with ice cold DMEM medium. To recover the fibers, the mixture was passed through a 70 µm strainer. Fibers were collected from the strainer, washed with 1x PBS and resuspended in RTL lysis buffer (for sequencing) or tripleXtractor reagent (for RNA extraction and qPCR). Samples in lysis buffer were sent to the Genomics Facility of Instituto Gulbenkian de Ciência (IGC) to proceed with library preparation and next generation sequencing. Briefly, SMART-Seq2 protocol was used to generate full-length cDNAs and sequencing libraries directly from lysed muscle fibers. The Nextera library preparation protocol (Nextera XT DNA Library Preparation kit, Illumina) was used for library preparation, including cDNA ‘tagmentation’, PCR-mediated adaptor addition, and library amplification. The libraries were sequenced using the NextSeq 2000 P2 Reagents (100 cycles) (Illumina). Sequencing data was extracted in FastQ format, resulting on average in approximately 30×10^6^ reads per sample and analyzed using Galaxy Europe platform^76,77^ and following a previously reported workflow^78^. The quality of FastQ reads was analyzed *FastQC*, which were then trimmed and filtered using *Cutadapt* and finally aligned against the mouse reference genome GRCm39 using *RNA STAR*. Read summarization was performed by assigning uniquely mapped reads to genomic features using *FeatureCounts* and then differential gene expression analysis comparing wildtype and *dy^W^* muscle fibers was performed using DESeq2. Functional analysis was performed with gProfiler platform (https://biit.cs.ut.ee/gprofiler/page/citing), Bubble plot representation was performed using SRplot^79^.

ChIP-seq data for C2C12 Myoblast and Myocyte (H3K4me3 peaks) were obtained from ENCODE Project (https://www.encodeproject.org/help/citing-encode/). Peaks/Gene association was performed using ChIPseeker^81^.

Gene lists of GO:0030036, GO:0042692, GO:0006281, GO:0006979, GO:0007005, GO:0007049 were obtained from Mouse Genome Informatics Web Site (MGI) (https://www.informatics.jax.org/mgihome/other/citation.shtml). The Venn diagram analysis was done using the Platform Bioinformatics & Evolutionary Genomics (https://bioinformatics.psb.ugent.be/webtools/Venn/) and the heatmaps with the platform Morpheus (https://software.broadinstitute.org/morpheus/).

### Quantification and statistical analysis

Image J/Fiji software was used to analyze immunofluorescence images and for densitometry analysis of westerns blots. Statistical analysis was conducted using GraphPad Prism 9 software. All distributed data are displayed as means ± standard deviation of the mean (SD) unless otherwise noted. Measurements between two groups were performed with Unpaired t-test. Groups of three or more were analyzed by ordinary one-way ANOVA with Dunnett’s multiple comparison test and two-way ANOVA with Tukey’s multiple comparisons test, depending on the number of variables analyzed. Statistical parameters for each experiment can be found within the corresponding figure legend. Serial cloner 2.6.1, SnapGene software and Synthego Performance Analysis, ICE Analysis (2019.v3.0) were used for sequencing analysis.

### Reagents and antibodies

All reagents, antibodies and resources are described on Key Resources Table.

### Key Resources Table

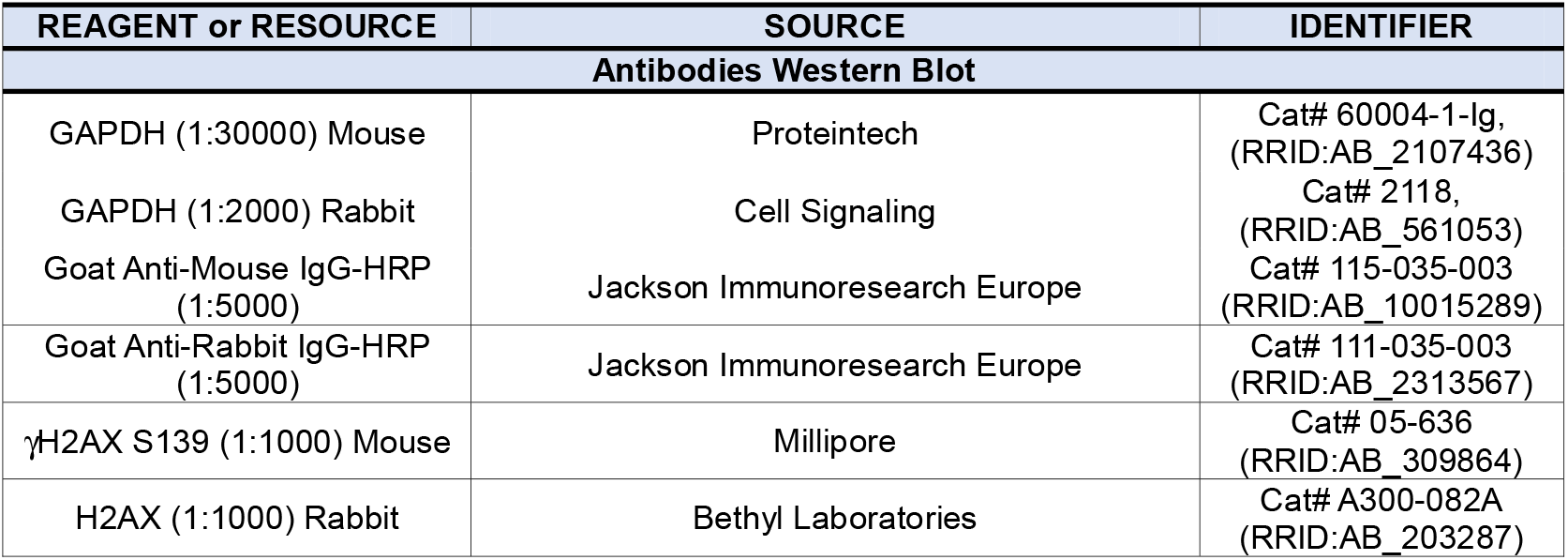
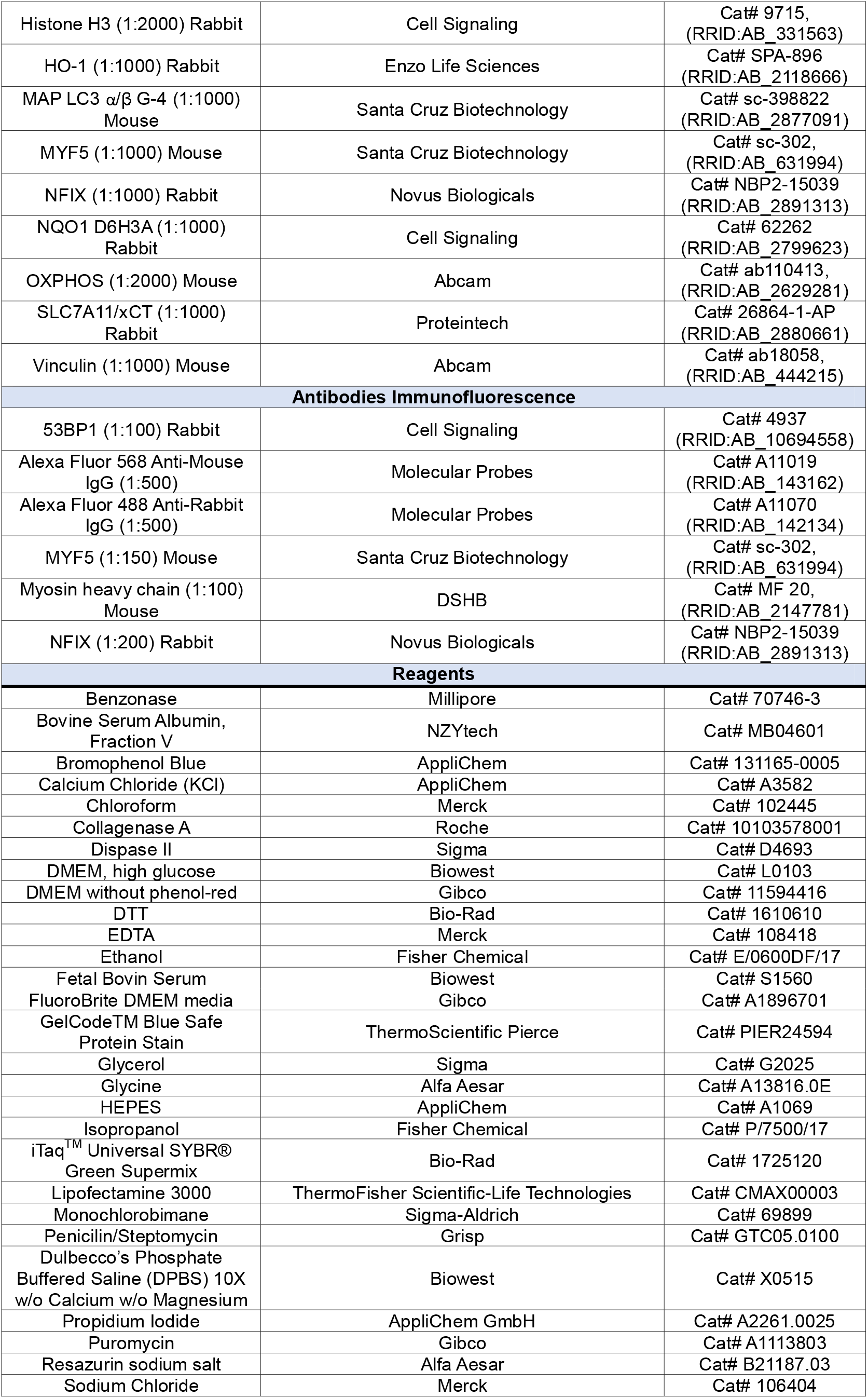
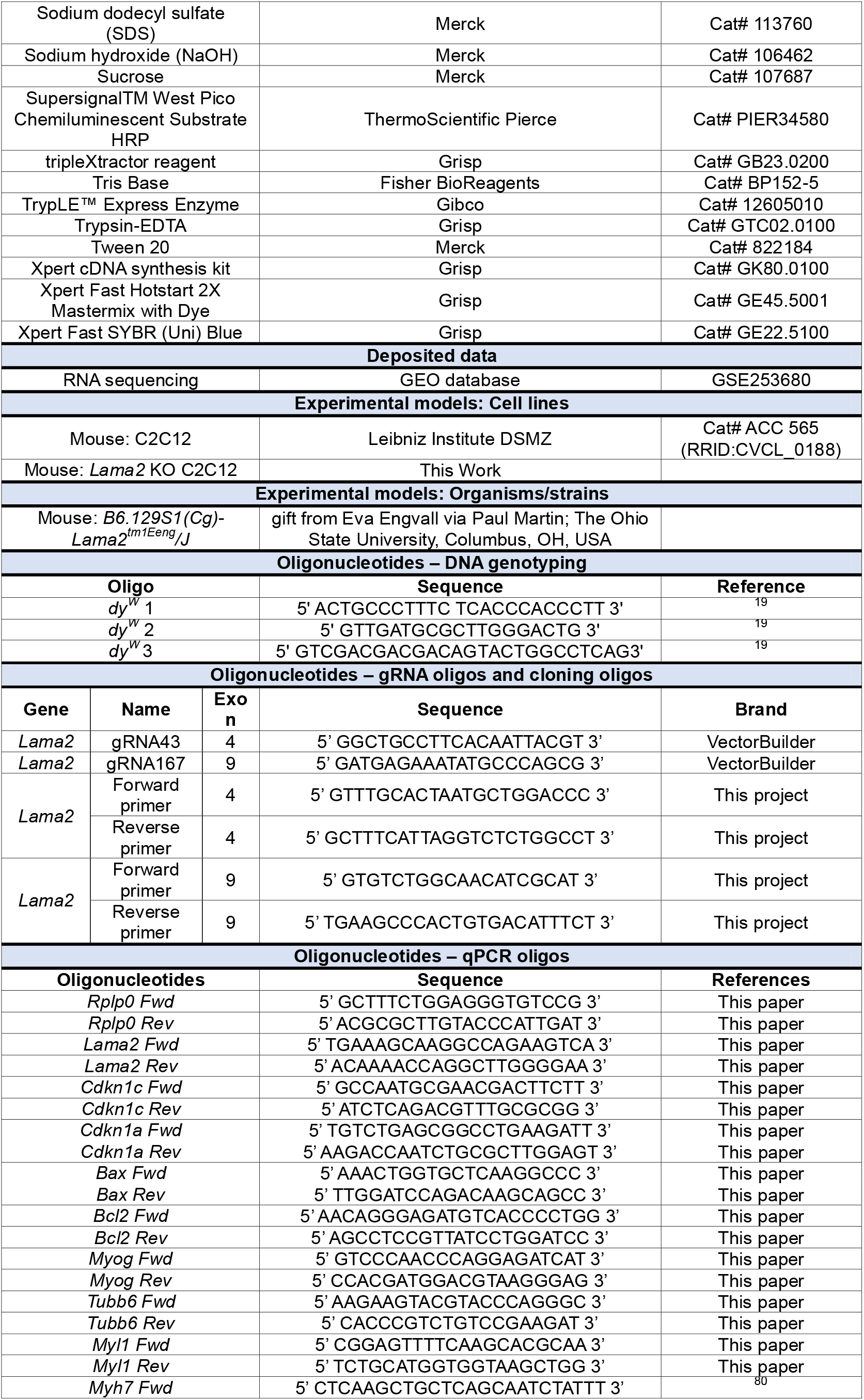
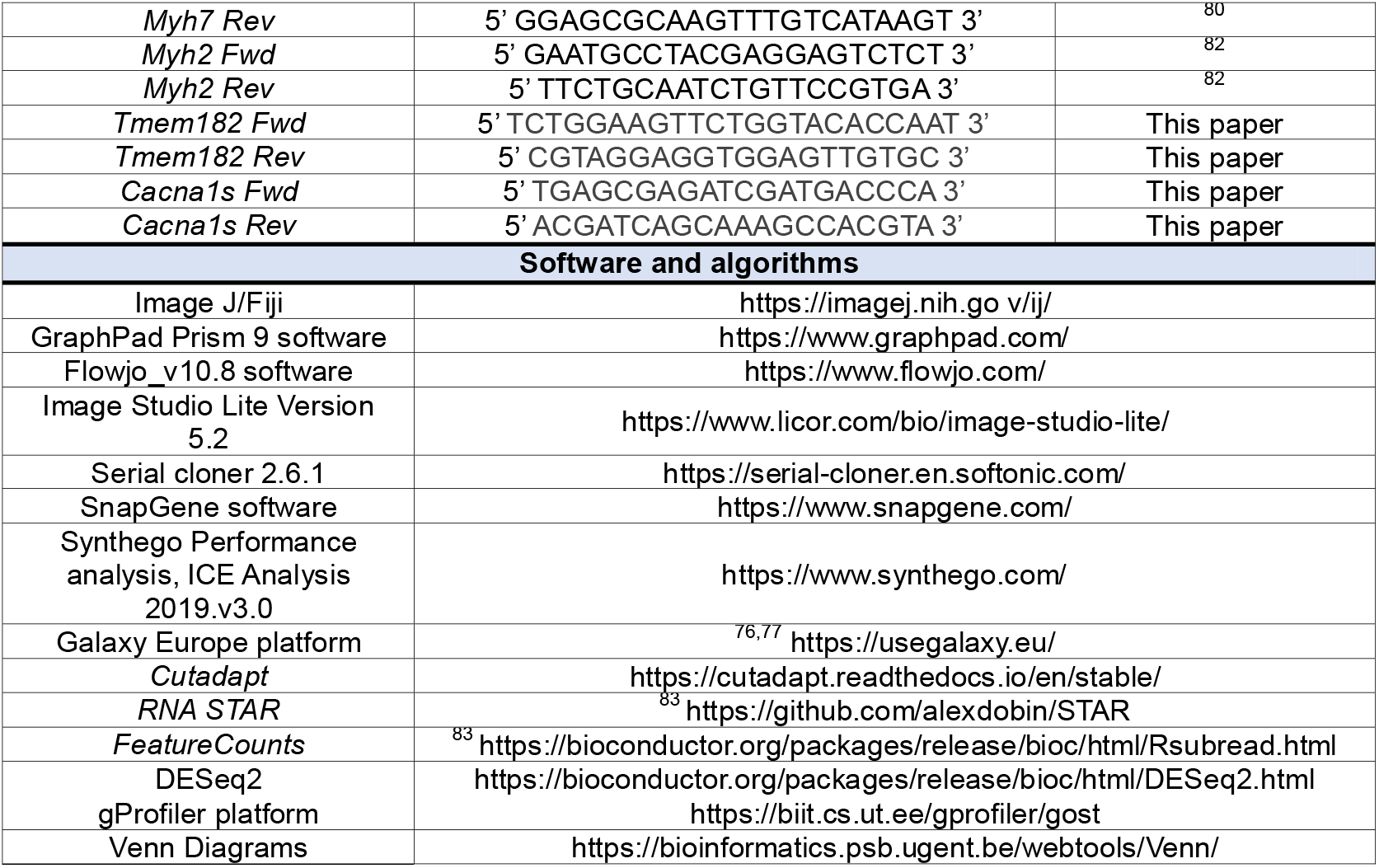

## Notes

### Competing Interest Statement

The authors have declared no competing interest.

