## Supplementary data for "Deregulation of multiple mechanisms shapes the onset of *LAMA2*-congenital muscular dystrophy"

**Supplementary Table 1.** List of the 191 genes with active expression based on H3K4me3 marker in C2C12 myotubes but downregulated in the RNAseq analysis of muscle fibers (described in Figure 4D).

| Gene Symbol | Gene Symbol | Gene Symbol | Gene Symbol | Gene Symbol |
|---|---|---|---|---|
| 1700008O03Rik | Cabp1 | Gm773 | Ncf2 | Slc6a4 |
| 1700017G19Rik | Cacna1s | Gm7854 | Nefl | Spats1 |
| 1700061J23Rik | Car5b | Gm960 | Neil1 | Tbx4 |
| 1700112H15Rik | Car6 | Gpr137c | Nkx2-2os | Tcf21 |
| 1700112J16Rik | Casz1 | Gpr156 | Nlrx1 | Tcte2 |
| 2610027K06Rik | Catsperg1 | Grhl1 | Nod2 | Tigd5 |
| 2610035F20Rik | Ccdc172 | Grik5 | Nog | Tmem154 |
| 2900005J15Rik | Ccdc87 | H2-T10 | Nrap | Tmem174 |
| 4632428C04Rik | Ccdc88b | Hapln4 | Nudt15 | Tmem182 |
| 4921525O09Rik | Cdkl4 | Hey2 | Nyx | Tmem74 |
| 4930405A10Rik | Cdsn | Hid1 | Obscn | Trim54 |
| 4930435F18Rik | Cercam | Hmcn2 | Odf4 | Trp63 |
| 4930449E01Rik | Ces5a | Hsf4 | P2rx3 | Tsks |
| 4930558J18Rik | Chrng | Hunk | Pax6 | Tspan32 |
| 4930565D16Rik | Cntd1 | Ifit1 | Pdzd7 | Ttc16 |
| 4930583K01Rik | Crb2 | Il11 | Phldb3 | Ttll13 |
| 4933406I18Rik | Crppa | Iqank1 | Phtf1os | Usp17la |
| 5430421F17Rik | Cstdc2 | Isl1 | Platr14 | Vinac1 |
| 6430550D23Rik | Dbndd1 | Islr2 | Platr26 | Vmn1r35 |
| 9230102K24Rik | Efcab1 | Kcnip3 | Plpp7 | Wfikkn2 |
| 9330104G04Rik | Efhb | Kcnj12 | Ppfia4 | Xirp1 |
| 9330111N05Rik | Efhd1 | Kif6 | Prdm8 | Zbtb49 |
| 9430014N10Rik | Eomes | Klhdc8b | Proser2 | Zfp493 |
| A830009L08Rik | Esrrb | Klhl14 | Ptpn5 | Zfp536 |
| Acsbg1 | Extl1 | Klhl33 | Rab3c | Zfp764 |
| Agap2 | F2 | Lncenc1 | Reep6 | Zfp831 |
| Alpk2 | Fbxl21 | Lrrc61 | Rgs9 | Zfp940 |
| Alpk3 | Fbxo16 | Ly86 | Rmst | Zfp994 |
| Alx1 | Fsip1 | Lynx1 | Sarm1 | Zim3 |
| Amz1 | Galnt15 | Map3k13 | Scn5a | Zkscan7 |
| Angptl6 | Gbx1 | Marveld3 | Sf3a2 | Zpbp2 |
| Ankrd23 | Glrp1 | Mdga1 | Sgip1 | |
| Ankrd55 | Gm10548 | Meig1 | Sgsm1 | |
| Apol10b | Gm15413 | Mfsd7a | Slc16a14 | |
| Arhgap8 | Gm15850 | Mlph | Slc16a3 | |
| Arhgef37 | Gm16894 | Mogat1 | Slc24a2 | |
| Bach2os | Gm17308 | Mucl3 | Slc25a23 | |
| BC051019 | Gm26760 | Mx2 | Slc2a4 | |
| Bicdl1 | Gm29688 | Myb | Slc2a6 | |
| Brinp3 | Gm5464 | Mypn | Slc4a5 | |

**Supplementary Table 2.** List of genes obtained from the Venn diagram analysis comparing the differentially expressed genes (DEGs) (p-value 0.05, log2 fold change +/-1.5) of wildtype vs. *dy*$^{W}$ muscle fibers (in Figure 4) with gene ontology analysis using the GO:0030036 Actin Cytoskeleton Organization.

| Actin Cytoskeleton Organization | | | | | | | | | | | |
|---|---|---|---|---|---|---|---|---|---|---|---|
| Downregulated | | | | | | | | Upregulated | | | |
| Gene symbol | Log 2 (FC) | Gene symbol | Log2 (FC) | Gene symbol | Log2 (FC) | Gene symbol | Log2 (FC) | Gene symbol | Log 2 (FC) | Gene symbol | Log2 (FC) |
| Actg1 | -2,41 | Dner | -4,55 | Mylk3 | -8,97 | Smyd3 | -1,56 | Asf1a | 3,41 | Qki | 2,24 |
| Adgrb1 | -3,99 | Dock2 | -4,68 | Myorg | -4,02 | T | -7,30 | Bmpr2 | 2,09 | Rgs2 | 1,84 |
| Adra1b | -9,49 | Flt3l | -5,23 | Mypn | -2,41 | Tbx1 | -7,58 | Bnip2 | 1,85 | Tomm70a | 1,55 |
| Akap6 | -1,89 | G6pd2 | -7,43 | Naglu | -3,56 | Tbx5 | -7,49 | Ccn4 | 2,93 | Yy1 | 1,91 |
| Alpk2 | -4,82 | Gata4 | -8,34 | Neu2 | -4,32 | Tcf23 | -3,94 | Cfh | 3,40 | Zeb1 | 1,69 |
| Alpk3 | -5,02 | Gata6 | -4,51 | Nfatc2 | -2,88 | Tmem182 | -7,64 | Ctnnb1 | 1,82 | | |
| Ankrd23 | -6,06 | Hey2 | -7,86 | Nln | -2,84 | Tnfsf14 | -7,61 | Efnb2 | 2,26 | | |
| Asb2 | -5,66 | Isl1 | -7,99 | Nphs1 | -4,56 | Trim72 | -3,91 | Hes1 | 1,59 | | |
| B4galnt2 | -8,72 | Krt19 | -6,53 | Nrap | -9,72 | Ttn | -2,35 | Hnrnpu | 2,14 | | |
| Barx2 | -7,43 | Lama1 | -3,18 | P2rx2 | -6,80 | Wfikkn1 | -3,84 | Ifrd1 | 2,63 | | |
| Bhlha15 | -6,44 | Lmod1 | -7,52 | Pi16 | -4,38 | Wfikkn2 | -8,35 | Igf1 | 2,83 | | |
| Bmp10 | -7,30 | Lrrc27 | -3,03 | Pin1rt1 | -7,60 | Xirp1 | -7,98 | Igfbp5 | 3,03 | | |
| C3 | -3,94 | Msx1 | -6,15 | Pld3 | -4,12 | Xk | -4,26 | Lox | 3,47 | | |
| Cacna1s | -8,57 | Mybpc3 | -8,98 | Plpp7 | -8,14 | Zfp418 | -10,05 | Nr3c1 | 2,01 | | |
| Cd53 | -4,35 | Myh6 | -9,49 | Popdc2 | -5,65 | | | Pdgfra | 1,88 | | |
| Cntnap2 | -2,56 | Myh7 | -7,91 | Prdm6 | -7,47 | | | Ppp3ca | 2,33 | | |
| Cxcl9 | -7,74 | Myhas | -7,47 | Rpl3l | -7,49 | | | Ppp3cb | 2,06 | | |

**Supplementary Table 3.** List of genes obtained from the Venn diagram analysis comparing the differentially expressed genes (DEGs) (p-value 0.05, log2 fold change +/-1.5) of wildtype vs. *dy*$^{W}$ muscle fibers (in Figure 4) with gene ontology analysis using the GO:0042692 Muscle Cell Differentiation

| Muscle Cell Differentiation | | | | | | | | | | | |
|---|---|---|---|---|---|---|---|---|---|---|---|
| Downregulated | | | | | | | | | | Upregulated | |
| Gene symbol | Log2 (FC) | Gene symbol | Log2 (FC) | Gene symbol | Log2 (FC) | Gene symbol | Log2 (FC) | Gene symbol | Log2 (FC) | Gene symbol | Log2 (FC) |
| Actb | -2,19 | Crhr2 | -8,26 | Insrr | -8,67 | Nlrp5 | -6,43 | Scin | -3,90 | Actr2 | 1,52 |
| Actg1 | -2,41 | Cryaa | -6,96 | Kash5 | -5,11 | Nphs1 | -4,56 | Serpinf2 | -5,49 | Antxr1 | 1,65 |
| Adgrb1 | -3,99 | Dmtn | -4,16 | Kbtbd13 | -5,22 | Nrap | -9,72 | Sh2b2 | -4,40 | Cald1 | 1,86 |
| Agap2 | -4,86 | Dnai3 | -3,99 | Krt19 | -6,53 | Ooep | -3,54 | Shank1 | -4,12 | Capza2 | 2,31 |
| Aif1l | -2,09 | Dock2 | -4,68 | Limk1 | -3,00 | Parvg | -8,61 | Shroom3 | -3,35 | Cul3 | 1,89 |
| Alox15 | -7,26 | Elmo3 | -2,87 | Lmod1 | -7,52 | Pawr | -8,91 | Specc1 | -2,12 | Epb41l2 | 1,74 |
| Ankrd23 | -6,06 | Ermn | -6,89 | Mlst8 | -5,28 | Pfn4 | -8,10 | Spire2 | -7,53 | Fmnl2 | 2,07 |
| Aqp2 | -8,17 | Espn | -8,74 | Msrb1 | -3,10 | Phactr3 | -7,69 | Sptb | -2,43 | Gja1 | 2,34 |
| Arhgap40 | -9,09 | Espnl | -8,29 | Mybpc3 | -8,98 | Pick1 | -1,95 | Sptbn5 | -8,23 | Marcks | 3,41 |
| Arhgap44 | -3,54 | Fam107a | -4,51 | Myh14 | -3,11 | Pls1 | -8,79 | Srcin1 | -3,99 | Myo1b | 1,67 |
| Asb2 | -5,66 | Fchsd1 | -5,58 | Myh6 | -9,49 | Ppargc1b | -4,46 | Tac1 | -8,33 | Nck1 | 2,55 |
| Bmp10 | -7,30 | Fhdc1 | -5,57 | Myh7 | -7,91 | Prkcq | -3,90 | Tacr1 | -8,30 | Nckap1 | 1,88 |
| Bst1 | -8,85 | Fmnl1 | -2,89 | Mylk3 | -8,97 | Pstpip2 | -4,89 | Trpm2 | -8,40 | Pdgfra | 1,88 |
| Capn10 | -3,01 | Frmd5 | -4,45 | Myo15a | -4,10 | Pxn | -3,94 | Ttn | -2,35 | Phactr2 | 2,37 |
| Carmil2 | -8,71 | Frmd7 | -6,79 | Myo1a | -8,11 | Rhoh | -10,38 | Ush1c | -6,05 | Pls3 | 2,60 |
| Carmil3 | -5,28 | Fzd10 | -9,16 | Myo3b | -7,00 | Rhpn1 | -9,34 | Vil1 | -3,57 | Rdx | 1,66 |
| Cdc42bpb | -2,55 | Ghrl | -5,94 | Myo5b | -4,63 | Rnd1 | -6,29 | Wasf3 | -2,81 | Rock2 | 1,58 |
| Cln3 | -2,47 | Ghsr | -7,51 | Myo5c | -7,00 | Rnd2 | -3,22 | Wipf3 | -5,80 | Sptbn1 | 1,93 |
| Clrn1 | -7,84 | Grhl3 | -7,82 | Myo7a | -2,35 | Rtkn | -3,55 | Xirp1 | -7,98 | Tgfbr1 | 1,95 |
| Coro7 | -2,84 | Hip1r | -4,92 | Myo7b | -4,23 | S1pr2 | -2,30 | | | Twf1 | 2,41 |
| Cpne6 | -9,17 | Hrg | -5,79 | Mypn | -2,41 | Samd14 | -3,42 | | | Zeb2 | 1,73 |

**Supplementary Table 4.** List of genes obtained from the Venn diagram analysis comparing the differentially expressed genes (DEGs) (p-value 0.05, log2 fold change +/-1.5) of wildtype vs. *dy*$^{W}$ muscle fibers (in Figure 4) with gene ontology analysis using the GO:0007049 Cell Cycle.

| Cell Cycle | | | | | | | | | | | |
|---|---|---|---|---|---|---|---|---|---|---|---|
| Downregulated | | | | | | | | Upregulated | | | |
| Gene symbol | Log2 (FC) | Gene symbol | Log2 (FC) | Gene symbol | Log2 (FC) | Gene symbol | Log2 (FC) | Gene symbol | Log2 (FC) | Gene symbol | Log2 (FC) |
| 1700028K03Rik | -7,72 | E4f1 | -3,17 | Misp | -4,97 | Spata17 | -7,57 | Actr2 | 1,52 | Nipbl | 1,63 |
| 4933427D14Rik | -4,31 | Ecrg4 | -4,38 | Morc2b | -7,28 | Spdya | -7,87 | Adamts1 | 1,99 | Orc4 | 1,91 |
| Actb | -2,19 | Enkd1 | -4,07 | Mov10l1 | -4,89 | Spdye4a | -7,69 | Angel2 | 1,74 | Pcna | 2,18 |
| Actl6b | -8,90 | Espl1 | -2,89 | Mre11a | -2,95 | Spire2 | -7,53 | Apc | 1,83 | Pds5b | 2,37 |
| Actr5 | -3,00 | Exo1 | -3,23 | Mrgprb2 | -6,87 | Spo11 | -7,27 | Arid2 | 1,64 | Phf10 | 2,09 |
| Ajuba | -2,77 | Fam107a | -4,51 | Msx1 | -6,15 | Sstr5 | -8,10 | Atrx | 1,64 | Pkd2 | 2,14 |
| Alox8 | -6,04 | Fkbp6 | -7,72 | Myb | -3,78 | Stag3 | -2,87 | Bnip2 | 1,85 | Plk2 | 1,60 |
| Ankfn1 | -9,28 | Flt3l | -5,23 | Nlrp5 | -6,43 | Stk33 | -7,79 | Btg1 | 2,09 | Ppp1r12a | 1,84 |
| Ankrd31 | -3,41 | Fsd1 | -8,35 | Nr2e1 | -3,49 | Stox1 | -8,47 | Ccnb2 | 2,12 | Ppp2ca | 1,91 |
| Ankrd53 | -5,87 | Fzd9 | -8,06 | Nudt15 | -6,77 | Syce1l | -8,02 | Ccng2 | 2,01 | Ppp2cb | 2,31 |
| Apbb1 | -4,87 | Gata3 | -3,86 | Nuf2 | -2,25 | Sycp2l | -8,11 | Ccni | 1,60 | Ppp2r2d | 1,58 |
| Aurkc | -8,09 | Gata4 | -8,34 | Ooep | -3,54 | Taf6 | -1,99 | Ccnl1 | 2,01 | Ppp3ca | 2,33 |
| Bid | -4,59 | Gata6 | -4,51 | Ovol1 | -7,67 | Tas2r121 | -7,28 | Cdc123 | 2,15 | Prpf40a | 1,58 |
| Birc7 | -7,80 | Gen1 | -2,47 | Pard6b | -5,60 | Tbx1 | -7,58 | Cdc27 | 1,90 | Pum2 | 1,94 |
| Brca2 | -3,41 | Gjc2 | -7,46 | Parp9 | -3,29 | Tert | -8,45 | Cdc73 | 1,67 | Rad21 | 2,23 |
| Brinp2 | -4,94 | Gm20824 | -6,76 | Pax6 | -5,69 | Tesmin | -8,78 | Chmp5 | 1,89 | Rdx | 1,66 |
| Brinp3 | -4,56 | Gm4297 | -6,63 | Pcid2 | -4,04 | Tex12 | -7,83 | Ckap2 | 2,29 | Rgs2 | 1,84 |
| Brip1 | -2,44 | Gm5934 | -7,99 | Phgdh | -2,91 | Tex19.1 | -7,49 | Cltc | 1,88 | Rock2 | 1,58 |
| Brme1 | -8,05 | Gm773 | -7,44 | Piwil1 | -8,48 | Tex19.2 | -7,52 | Csnk1a1 | 1,60 | Rps6ka3 | 1,93 |
| Btbd18 | -9,99 | Gmnc | -7,72 | Piwil4 | -5,96 | Tex24 | -8,62 | Ctcf | 1,86 | Sbds | 1,72 |
| Btg1c | -7,01 | Gpr132 | -7,05 | Plk5 | -8,63 | Tgm1 | -4,89 | Ctnnb1 | 1,82 | Septin11 | 2,23 |
| Btn2a2 | -7,67 | Hepacam | -7,86 | Pml | -2,62 | Tjp3 | -3,24 | Cul3 | 1,89 | Skil | 3,10 |
| Camk2b | -3,00 | Hepacam2 | -5,87 | Prdm11 | -4,60 | Tm4sf5 | -5,41 | Dctn6 | 1,93 | Smarca5 | 1,52 |
| Ccnf | -2,37 | Hnf4a | -9,06 | Prdm9 | -3,35 | Tmem67 | -3,24 | Ddx3x | 2,82 | Smc2 | 1,94 |
| Cdk15 | -3,76 | Hyal1 | -4,57 | Prkcq | -3,90 | Trim36 | -4,35 | Dr1 | 1,58 | Smc3 | 1,79 |
| Cdk20 | -7,96 | Iho1 | -3,13 | Psma8 | -4,14 | Trim71 | -3,89 | Eid1 | 2,40 | Smc4 | 1,69 |
| Cdk5rap1 | -3,43 | Ing4 | -8,28 | Rab11fip3 | -2,29 | Trim75 | -7,47 | Eif4e | 1,92 | Son | 2,65 |
| Cenpo | -5,14 | Ins1 | -6,28 | Rad51c | -4,66 | Trp63 | -4,55 | Epb41l2 | 1,74 | Sptbn1 | 1,93 |
| Cenpt | -2,29 | Insm2 | -6,75 | Rcc1 | -1,59 | Trp73 | -3,95 | Fap | 2,46 | Stag2 | 1,72 |
| Clgn | -9,10 | Kash5 | -5,11 | Recql5 | -2,21 | Ttbk1 | -8,95 | Gja1 | 2,34 | Tardbp | 2,18 |
| Cntd1 | -7,09 | Kifc5b | -3,48 | Rgs14 | -9,14 | Tuba8 | -7,37 | Gnai3 | 2,08 | Tcim | 3,92 |
| Crnn | -6,18 | Klhdc8b | -4,97 | Rmi2 | -9,07 | Tubb4a | -8,45 | Gtf2b | 2,06 | Tlk2 | 1,85 |
| Crocc | -2,99 | L3mbtl1 | -4,79 | Rnf112 | -6,82 | Tube1 | -5,79 | Haus3 | 2,47 | Top2b | 1,87 |
| Ctc1 | -2,82 | Lep | -7,26 | Rnf8 | -2,78 | Tunar | -9,01 | Heca | 2,30 | Trim37 | 1,66 |
| Cts7 | -8,67 | Lif | -7,55 | Rsph1 | -3,81 | Ube2u | -4,00 | Hes1 | 1,59 | Tsc22d2 | 2,05 |
| Cul9 | -3,56 | Lig3 | -3,25 | Rtkn | -3,55 | Ush1c | -6,05 | Hnrnpu | 2,14 | Uchl5 | 2,51 |
| Cuzd1 | -7,36 | Lig4 | -5,17 | Rxfp3 | -8,32 | Usp26 | -7,96 | Igf1 | 2,83 | Ufl1 | 2,09 |
| Cyp27b1 | -6,83 | M1ap | -7,17 | Septin1 | -3,66 | Usp44 | -7,63 | Ik | 1,83 | Usp47 | 1,68 |
| D7Ertd443e | -9,74 | Madd | -3,81 | Sgo2b | -7,00 | Xrcc3 | -4,20 | Ing1 | 2,10 | Wapl | 1,71 |
| Ddx11 | -2,40 | Mael | -7,06 | Shcbp1l | -2,97 | Zbtb49 | -3,65 | Kif2a | 1,85 | Yy1 | 1,91 |
| Deup1 | -7,56 | Majin | -6,73 | Sipa1 | -2,41 | Zfp365 | -8,61 | Larp7 | 1,85 | Zfp207 | 1,54 |

| | | | | | | | | | | | |
|---|---|---|---|---|---|---|---|---|---|---|---|
| Dmrt1 | -5,93 | Map10 | -8,47 | Six3 | -4,77 | Zfp369 | -2,93 | Lrrcc1 | 2,25 | Zfp36l2 | 3,01 |
| Dmrtc2 | -6,65 | Map1s | -1,88 | Slc25a31 | -5,69 | | | Mbtd1 | 1,68 | | |
| Dnmt3l | -4,06 | Mapk15 | -8,51 | Slc26a8 | -3,70 | | | Mki67 | 1,56 | | |
| Dpf1 | -4,06 | Mdc1 | -2,11 | Slc6a4 | -5,82 | | | Nabp1 | 1,86 | | |
| E2f1 | -3,10 | Mei1 | -10,02 | Slfn1 | -7,61 | | | Nfia | 1,73 | | |
| E2f2 | -4,02 | Meig1 | -7,67 | Spag8 | -5,94 | | | Nfib | 2,10 | | |

**Supplementary Table 5.** List of genes obtained from the Venn diagram analysis comparing the differentially expressed genes (DEGs) (p-value 0.05, log2 fold change +/-1.5) of wildtype vs. *dy*$^{W}$ muscle fibers (in Figure 4) with gene ontology analysis using the GO:0006281 DNA Repair.

| DNA repair | | | | | | | | | |
|---|---|---|---|---|---|---|---|---|---|
| Downregulated | | | | | | Upregulated | | | |
| Gene symbol | Log2 (FC) | Gene symbol | Log2 (FC) | Gene symbol | Log2 (FC) | Gene symbol | Log2 (FC) | Gene symbol | Log2 (FC) |
| Actb | -2,19 | Hrob | -3,69 | Poln | -3,14 | Actr2 | 1,52 | Smc3 | 1,79 |
| Actl6b | -8,90 | Kash5 | -5,11 | Prdm9 | -3,35 | Arid2 | 1,64 | Smc4 | 1,69 |
| Actr5 | -3,00 | Kdm4d | -8,68 | Pwwp3a | -2,24 | Asf1a | 3,41 | Supt20 | 1,67 |
| Adprs | -2,62 | Lig3 | -3,25 | Rad51c | -4,66 | Atrx | 1,64 | Uchl5 | 2,51 |
| Apbb1 | -4,87 | Lig4 | -5,17 | Recql5 | -2,21 | Bod1l | 1,71 | Ufl1 | 2,09 |
| Ascc1 | -3,80 | Majin | -6,73 | Rmi2 | -9,07 | Eny2 | 2,09 | Usp47 | 1,68 |
| Brca2 | -3,41 | Mdc1 | -2,11 | Rnf8 | -2,78 | Gtf2h2 | 1,57 | Usp7 | 2,11 |
| Brip1 | -2,44 | Mgme1 | -3,11 | Spire2 | -7,53 | Kin | 2,48 | Xrn2 | 1,98 |
| Brme1 | -8,05 | Mgmt | -4,67 | Spo11 | -7,27 | Mbtd1 | 1,68 | Yy1 | 1,91 |
| Cyren | -3,86 | Morc2b | -7,28 | Supt7l | -3,30 | Nabp1 | 1,86 | | |
| Ddx11 | -2,40 | Mre11a | -2,95 | Taf6 | -1,99 | Nipbl | 1,63 | | |
| Dpf1 | -3,41 | Neil1 | -4,88 | Tdg-ps | -5,09 | Pcna | 2,18 | | |
| Endov | -6,17 | Neil2 | -9,34 | Tex12 | -7,83 | Pds5b | 2,37 | | |
| Exo1 | -3,23 | Ooep | -3,54 | Ttf2 | -3,54 | Phf10 | 2,09 | | |
| Eya2 | -8,60 | Parp9 | -3,29 | Ung | -4,01 | Psme4 | 2,00 | | |
| Fan1 | -9,80 | Pml | -2,62 | Xrcc3 | -4,20 | Rad21 | 2,23 | | |
| Gen1 | -2,47 | Pold2 | -1,91 | Zfp365 | -8,61 | Smarca5 | 1,52 | | |
| Ggn | -7,87 | Polh | -3,65 | | | Smc2 | 1,94 | | |

**Supplementary Table 6.** List of genes obtained from the Venn diagram analysis comparing the differentially expressed genes (DEGs) (p-value 0.05, log2 fold change +/-1.5) of wildtype vs. *dy*$^{W}$ muscle fibers (in Figure 4) with gene ontology analysis using the GO: 0006979 Response to Oxidative Stress.

| Oxidative Stress Response | | | | | | | | | |
|---|---|---|---|---|---|---|---|---|---|
| Downregulated | | | | | | | | Upregulated | |
| Gene symbol | Log2 (FC) | Gene symbol | Log2 (FC) | Gene symbol | Log2 (FC) | Gene symbol | Log2 (FC) | Gene symbol | Log2 (FC) |
| Adipoq | -7,22 | Keap1 | -2,50 | Nqo1 | -4,05 | Slc4a1 | -5,43 | Hsph1 | 1,92 |
| Adprs | -2,62 | Lck | -2,98 | Nudt15 | -6,77 | Slc4a11 | -9,40 | Idh1 | 1,82 |
| Aldh3b1 | -8,20 | Lcn2 | -7,74 | Pawr | -8,91 | Sod3 | -4,29 | Pdgfra | 1,88 |
| Alox5 | -4,55 | Lpo | -6,76 | Pax2 | -7,14 | Stc2 | -5,98 | Ppp2cb | 2,31 |
| Cryaa | -6,96 | Hnf1a | -8,99 | Pex10 | -4,97 | Stox1 | -8,47 | Prdx3 | 1,82 |
| Epor | -6,10 | Meak7 | -5,49 | Pml | -2,62 | Tacr1 | -8,30 | Psip1 | 2,09 |
| Epx | -7,41 | Mmp3 | -3,31 | Ppargc1b | -4,46 | Tat | -6,90 | Sod1 | 2,46 |
| G6pd2 | -7,43 | Mpv17l | -6,58 | Ptprn | -9,08 | Thg1l | -5,23 | | |
| Gpx2 | -8,50 | Mtf1 | -3,05 | Pyroxd1 | -3,90 | Tlr4 | -6,58 | | |
| Gpx5 | -8,16 | Naglu | -3,56 | Rbm11 | -7,73 | Trpm2 | -8,40 | | |
| Gpx6 | -7,68 | Neil1 | -4,88 | Reg3b | -7,57 | Ucp3 | -8,08 | | |
| Hk3 | -4,05 | Ngb | -7,51 | Rgs14 | -9,14 | Wnt16 | -5,51 | | |
| Hyal1 | -4,57 | Nme8 | -8,41 | Rnf112 | -6,82 | | | | |

**Supplementary Table 7.** List of genes obtained from the Venn diagram analysis comparing the differentially expressed genes (DEGs) (p-value 0.05, log2 fold change +/-1.5) of wildtype vs. $dy^W$ muscle fibers (in Figure 4) with gene ontology analysis using the GO:0007005 Mitochondria Organization.

| Mitochondrion Organization | | | | | | | | | | | |
|---|---|---|---|---|---|---|---|---|---|---|---|
| Downregulated | | | | | | | | Upregulated | | | |
| Gene symbol | Log2 (FC) | Gene symbol | Log2 (FC) | Gene symbol | Log2 (FC) | Gene symbol | Log2 (FC) | Gene symbol | Log2 (FC) | Gene symbol | Log2 (FC) |
| 2610042 L04Rik | -5,37 | Dmac2 | -5,09 | Mgme1 | -3,11 | Slc25a33 | -2,25 | Atg3 | 2,09 | Ssbp1 | 2,42 |
| Adck1 | -1,89 | Dnaja3 | -2,10 | Miga2 | -3,55 | Slc4a5 | -8,61 | Cfh | 3,40 | Timm22 | 2,18 |
| Afg1l | -4,36 | Dnajc11 | -2,19 | Mpv17l | -6,58 | Spata18 | -7,14 | Dctn6 | 1,93 | Tomm20 | 1,91 |
| Afg3l1 | -1,78 | Eya2 | -8,60 | Myh14 | -3,11 | Stox1 | -8,47 | Hsp90aa1 | 2,30 | Tomm70a | 1,55 |
| Agtpbp1 | -1,79 | Ggnbp1 | -6,42 | Nod2 | -6,10 | Stpg1 | -7,94 | Igf1 | 2,83 | Tug1 | 1,79 |
| Atcay | -5,65 | Hap1 | -9,01 | Nptx1 | -4,00 | Tert | -8,45 | Man2a1 | 2,29 | Vps54 | 1,97 |
| Atg4d | -8,10 | Hip1r | -4,92 | P2rx7 | -4,70 | Thg1l | -5,23 | Marcks | 3,41 | Yme1l1 | 1,62 |
| Atg9b | -8,88 | Hrk | -7,16 | Pisd | -2,10 | Tnfsf10 | -5,36 | Mcl1 | 1,98 | Zdhhc6 | 2,11 |
| Bhlha15 | -6,44 | Irgm2 | -7,97 | Pla2g6 | -4,20 | Tomm40 | -2,41 | Ogt | 2,34 | | |
| Bid | -4,59 | Kif28 | -7,83 | Ptpn5 | -9,38 | Trp73 | -3,95 | Ppp2cb | 2,31 | | |
| Capn10 | -3,01 | Letm1 | -2,45 | Rcc1l | -3,27 | Tymp | -7,90 | Prdx3 | 1,82 | | |
| Cntnap2 | -2,56 | Lig3 | -3,25 | Sco1 | -4,50 | | | Pum2 | 1,94 | | |
| Cryaa | -6,96 | Lncbate1 | -5,92 | Siah3 | -6,41 | | | Ralbp1 | 1,89 | | |
| Dhodh | -3,60 | Lyrm7 | -5,00 | Slc25a31 | -5,69 | | | Snx7 | 1,81 | | |

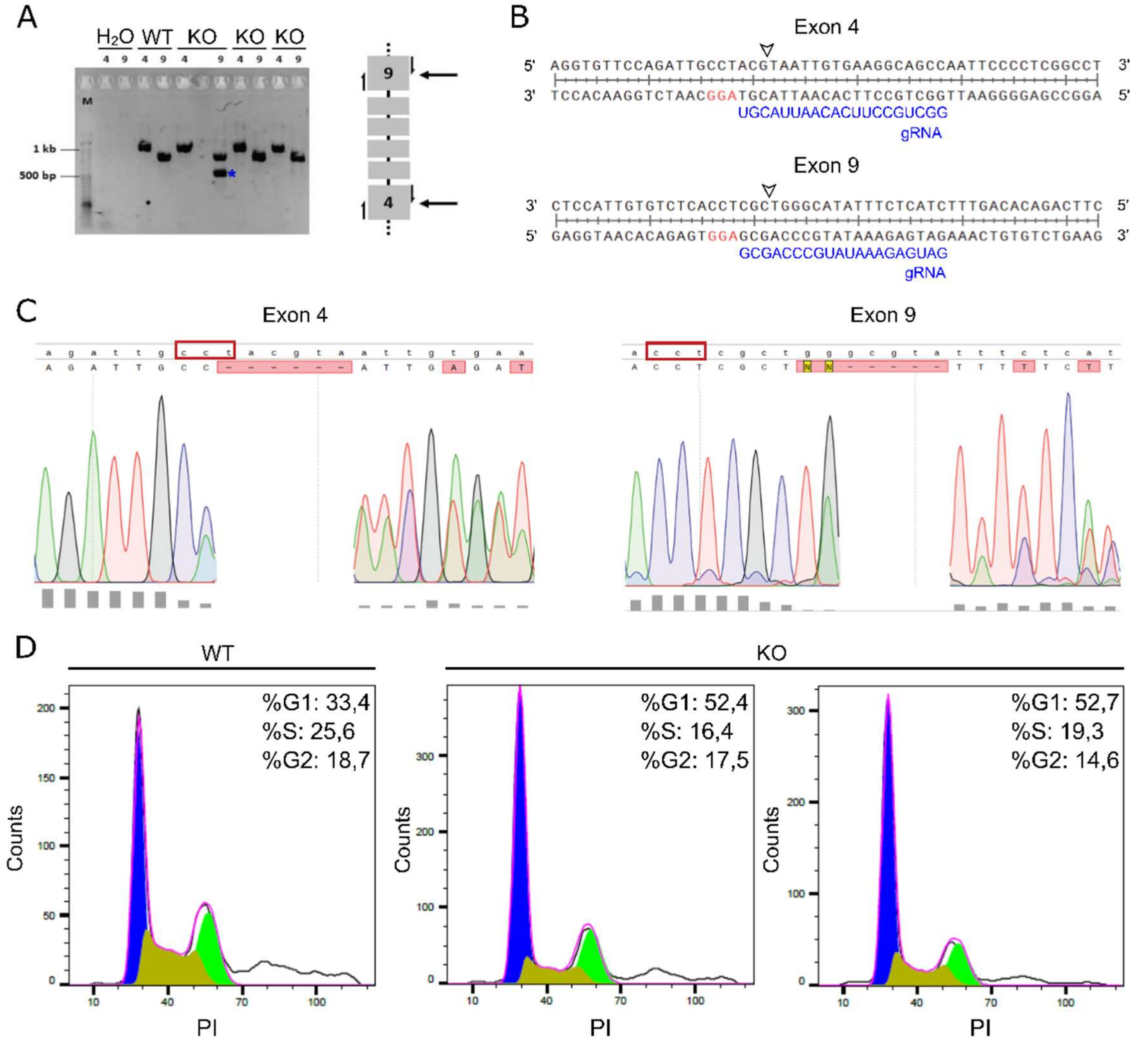


**Supplementary Figure 1 Characterization of the mutations on the generated *Lama2* knockout cell lines. A)** Genomic DNA from wildtype (WT) C2C12 cells and three independent *Lama2* knockout single cell clones (KO) was amplified through PCR using either the Lama2_exon4_Fwd and Lama2_exon4_Rev primers or the Lama2_exon9_Fwd and Lama2_exon9_Rev primers. *Lama2*-KO indicates three independent single cell clones. The negative control ($H_2O$) of the PCR is also shown. The blue asterisk indicates the deletion band. On the right, a diagram shows exons 4 and 9 of the *Lama2* gene targeted for the generation of the single cell clones. The half-arrows represent cut-site primers used to verify the presence of indels. **B)** Sequence comparison of the *Lama2* gene with gRNA sequences targeting the exons 4 (upper) and 9 (bottom). The gRNA and PAM sequences are shown in blue and red, respectively. The potential cleavage sites are indicated by arrowheads. **C)** Alignment of the WT sequences (lowercase) with the sequence tracers (uppercase) for the exons 4 (on the left) and 9 (on the right) of a *Lama2* KO single cell clone. The red boxes indicate the complementary sequences to the PAM sites and the bottom grey rectangles show the quality values of the chromatogram data. The primers Lama2_exon4_Fwd and Lama2_exon9_Rev were used for sequencing the cutting site on exon 4 and 9, respectively. The alignments were performed using the SnapGene software (www.snapgene.com). **D)** Representative histogram plot of flow cytometry cell cycle analysis (relative to Figure 1D). Percentages of cells in the different cell cycle phases are shown.

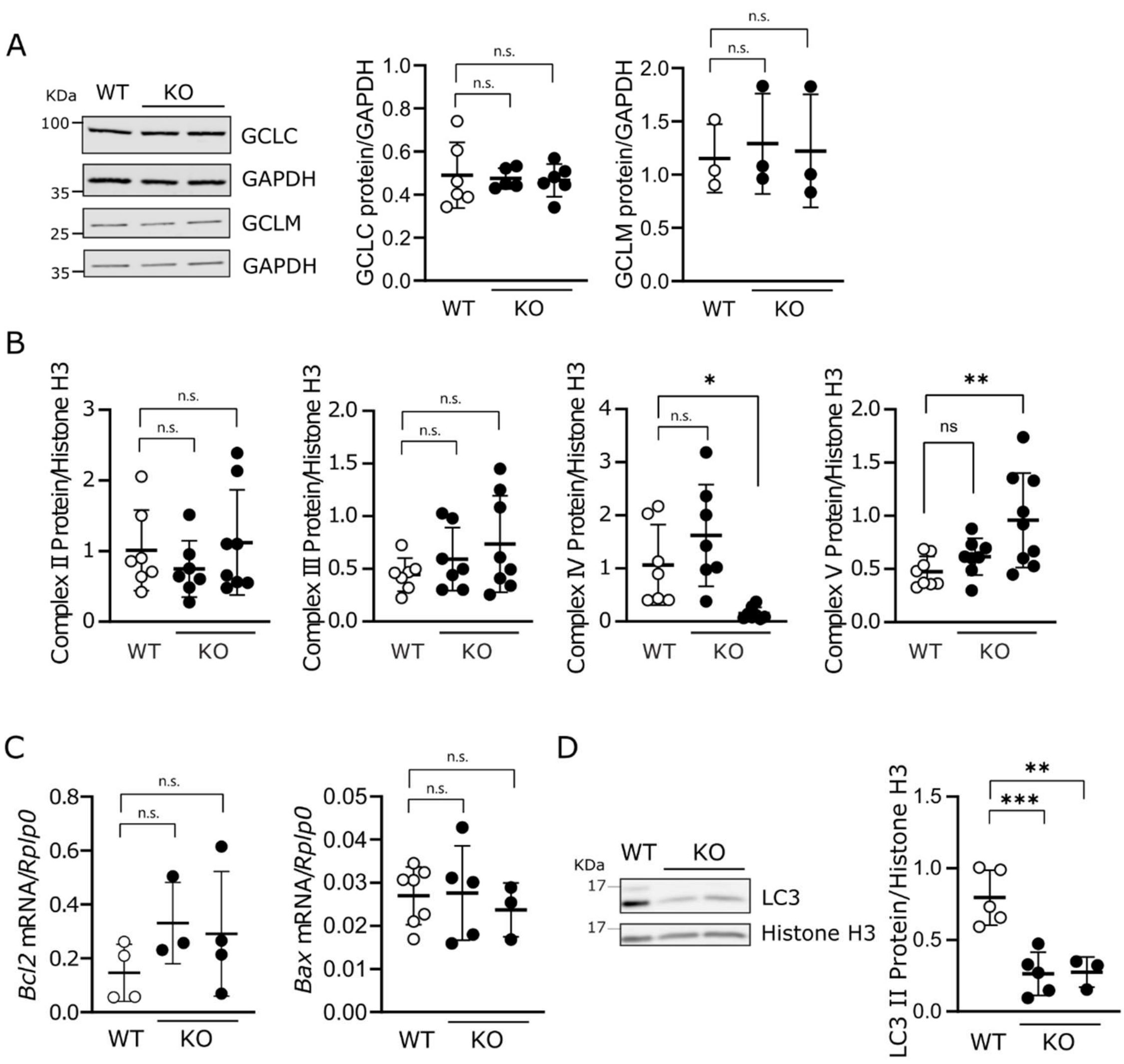


**Supplementary Figure 2 Glutathione pathway enzymes, mitochondrial complexes, apoptosis and autophagy were analyzed in Lama2-defficiency. A)** Wildtype (WT) C2C12 cells and two independent *Lama2* knockout single cell clones (KO) were harvested for western blot analysis with anti-GCLC and anti-GCLM antibodies (left panel). GAPDH was used as loading control. Two KO lanes represent experiments performed with two independent KO cell lines. n=3-6 samples per group collected independently. Densitometry analysis is shown on the right. **B)** Densitometry analysis of the western blot with anti-OXPHOS antibody shown on Figure 1 H. n=7-9 samples per group collected independently. **C)** WT and KO C2C12 cells were harvested, the RNA was extracted and the expression of *Bcl2* and *Bax* was analyzed by qPCR. n=3-7 independent samples per group collected independently. **D)** WT and KO C2C12 were harvested for western blot analysis with anti-LC3 antibody. Histone H3 was used as loading control. Two KO lanes represent experiments performed with KO cell lines. n=5-9 samples per group collected independently. Densitometry analysis of LC3_II is shown on the right. Statistical analysis was done using Ordinary One-way ANOVA by Dunnett's multiple comparisons test for A), B), C) and D). P-values: n.s. = not significant; * p<0.05; ** p<0.01.

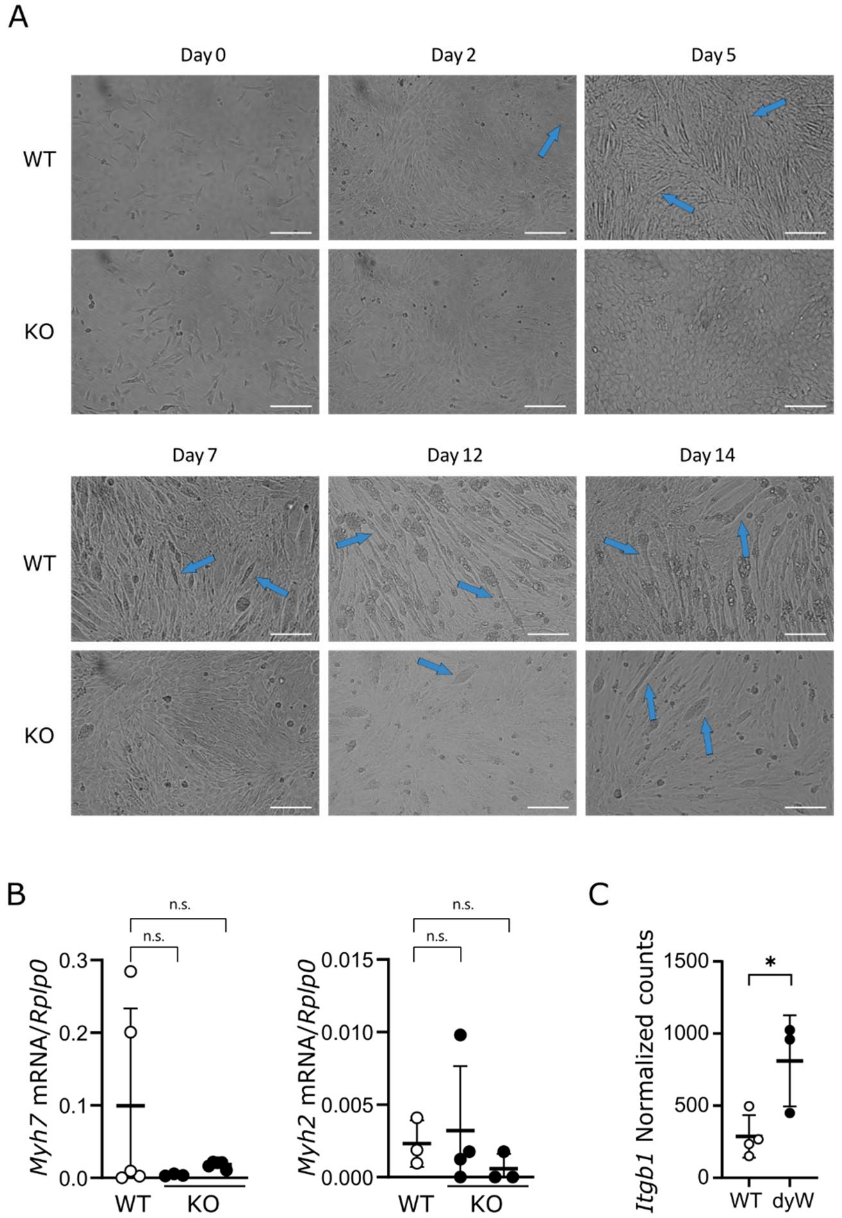


**Supplementary Figure 3 Differentiation is impaired in the absence of *Lama2* in 2D cultures.** **A)** Representative phase-contrast images of C2C12 wildtype (WT) cells and two independent *Lama2* knockout single cell clones (KO) cultured for 14 days in order to induce differentiation. Images were taken on days 0, 2, 5, 7, 12 and 14. Images of two independent *Lama2* KO clones were analyzed separately. Full blue arrows indicate myotubes. Scale bar: 200 μm. **B)** *Myh7* and *Myh2* were analyzed by qPCR in WT and KO C2C12 cells after 5 days in culture to induce differentiation. Expression levels were normalized using the housekeeping gene *Rplp0*. n=3-5 samples per group collected independently. **C)** Normalized counts of the gene *Itgb1*, selected from the analysis on Figure 4. n=3-4 independent samples per genotype. Statistical analysis performed using Ordinary One-way ANOVA for B) and Unpaired T-test for C). P-value: n.s.= not significant; * $p<0.05$.

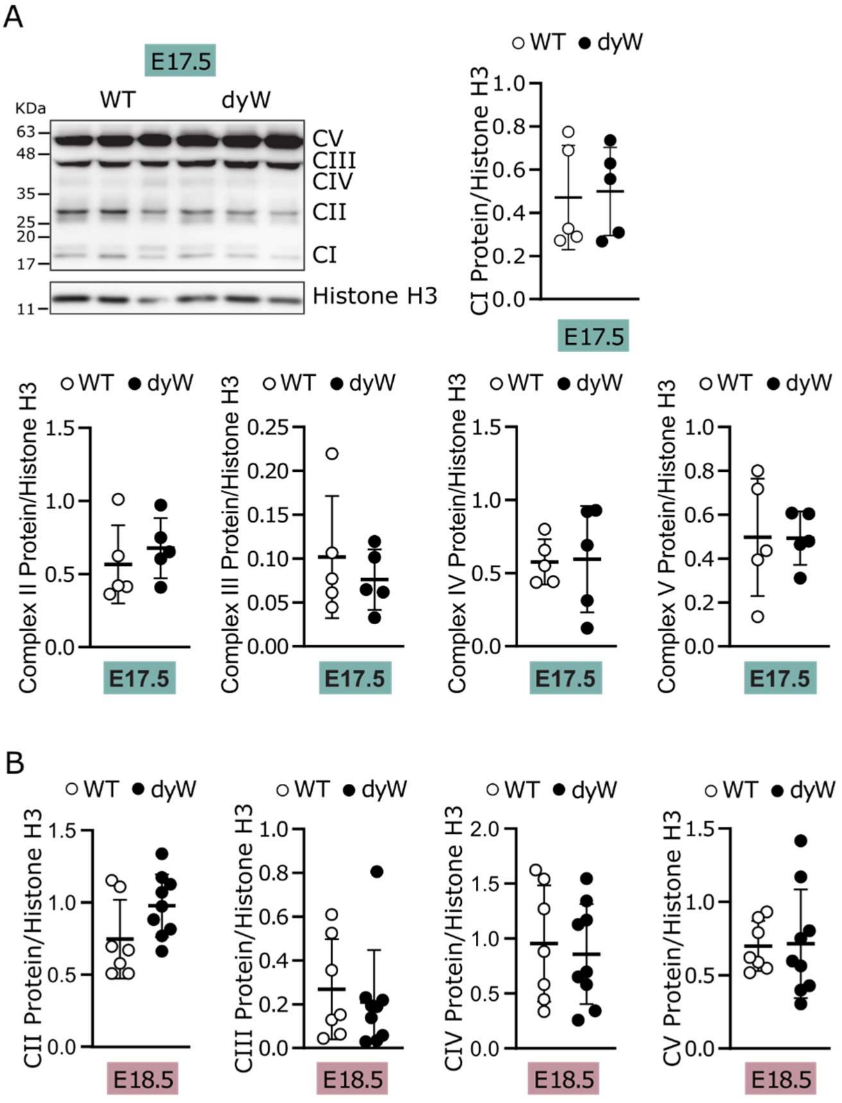


**Supplementary Figure 4 mitochondrial complexes analysis at E17.5 and E18.5. A)** Epaxial muscles were collected from wildtype and *dy*$^{W}$ fetuses at E17.5, the protein extracts were analyzed by western blot and the membranes probed with OXPHOS (CI - complex I; CII - complex II; CIII - complex III; CIV - complex IV; CV - complex V) antibodies. Histone H3 was used as loading control. Densitometry analysis of CI is shown on the right and densitometry analysis of CII, CIII, CIV and CV is shown below. n=5 fetuses per genotype. **B)** Densitometry analysis of the western blot with anti-OXPHOS (CII - complex II; CIII - complex III; CIV - complex IV; CV - complex V) antibody shown on Figure 7 C. n=7-9 fetuses per genotype. Statistical analysis was performed using Student's t-test for A) and B).
